# Morphogen gradient orchestrates pattern-preserving tissue morphogenesis via motility-driven (un)jamming

**DOI:** 10.1101/2022.05.16.492018

**Authors:** Diana Pinheiro, Roland Kardos, Édouard Hannezo, Carl-Philipp Heisenberg

## Abstract

Embryo development requires both biochemical signalling generating patterns of cell fates and active mechanical forces driving tissue shape changes. Yet, how these fundamental processes are coordinated in space and time, and, especially, how tissue patterning is preserved despite the complex cellular flows occurring during morphogenesis, remains poorly understood. Here, we show that a Nodal/TGF-β morphogen gradient orchestrates pattern-preserving mesendoderm internalization movements during zebrafish gastrulation by triggering a motility-driven (un)jamming transition. We find that graded Nodal signalling, in addition to its highly conserved role in mesendoderm patterning, mechanically subdivides the tissue into a small fraction of highly protrusive leader cells able to locally unjam and thus autonomously internalize, and less protrusive followers, which remain jammed and need to be pulled inwards by the leaders. Using minimal particle-based simulations and experimental perturbations, we further show that this binary mechanical switch, when combined with Nodal-dependent preferential adhesion coupling leaders to followers, is critical for triggering collective and orderly mesendoderm internalization, thus preserving tissue patterning. This provides a simple, yet quantitative, theoretical framework for how a morphogen-encoded (un)jamming transition can bidirectionally couple tissue mechanics with patterning during complex three-dimensional morphogenesis.

---

Embryogenesis involves precise and highly reproducible cell and tissue movements, which shape the embryo and position the newly specified progenitor cells along the future body axes. Experimental and theoretical work over the last decades has provided insight into the molecular signals and gene regulatory networks involved in tissue patterning. Many of these signals were shown to function as morphogens, diffusible molecules which form signalling gradients within tissues due to self-organized or pre-patterned cues, and induce cell fate in a concentration-dependent manner^1,2^. Likewise, considerable progress has been made in understanding the cellular processes and active mechanical forces involved in sculpting tissues and embryos^3–12^. However, while tissue patterning and morphogenesis must be tightly coordinated during development, the mechanochemical mechanisms underlying such coordination remain poorly understood.

Gastrulation is an excellent model to address this question, as it constitutes the first major morphogenetic event in development during which the different germ layers - ectoderm, mesoderm and endoderm - are both specified and shaped. During zebrafish gastrulation, the highly conserved class of Nodal/TGF-β morphogens form a signalling gradient at the blastoderm margin and induce mesoderm and endoderm (mesendoderm) cell fate specification in a dose-dependent manner (Fig. 1a)^13–17^. Since Nodal signals act primarily before gastrulation^18^, mesendoderm cell fate allocation is largely completed prior or concomitant with the onset of its internalization movements beneath the ectodermal layer^16,19^. Although, in multiple organisms, mesendoderm internalization involves various degrees of epithelial-to-mesenchymal transitions and large-scale cellular flows, the relative position of mesendoderm progenitors before internalization is indicative of their ultimate fate and position after gastrulation^16,19–21^. Yet, how such positional information is preserved during the complex three-dimensional (3D) flows associated with gastrulation remains unclear.

**Figure 1.**
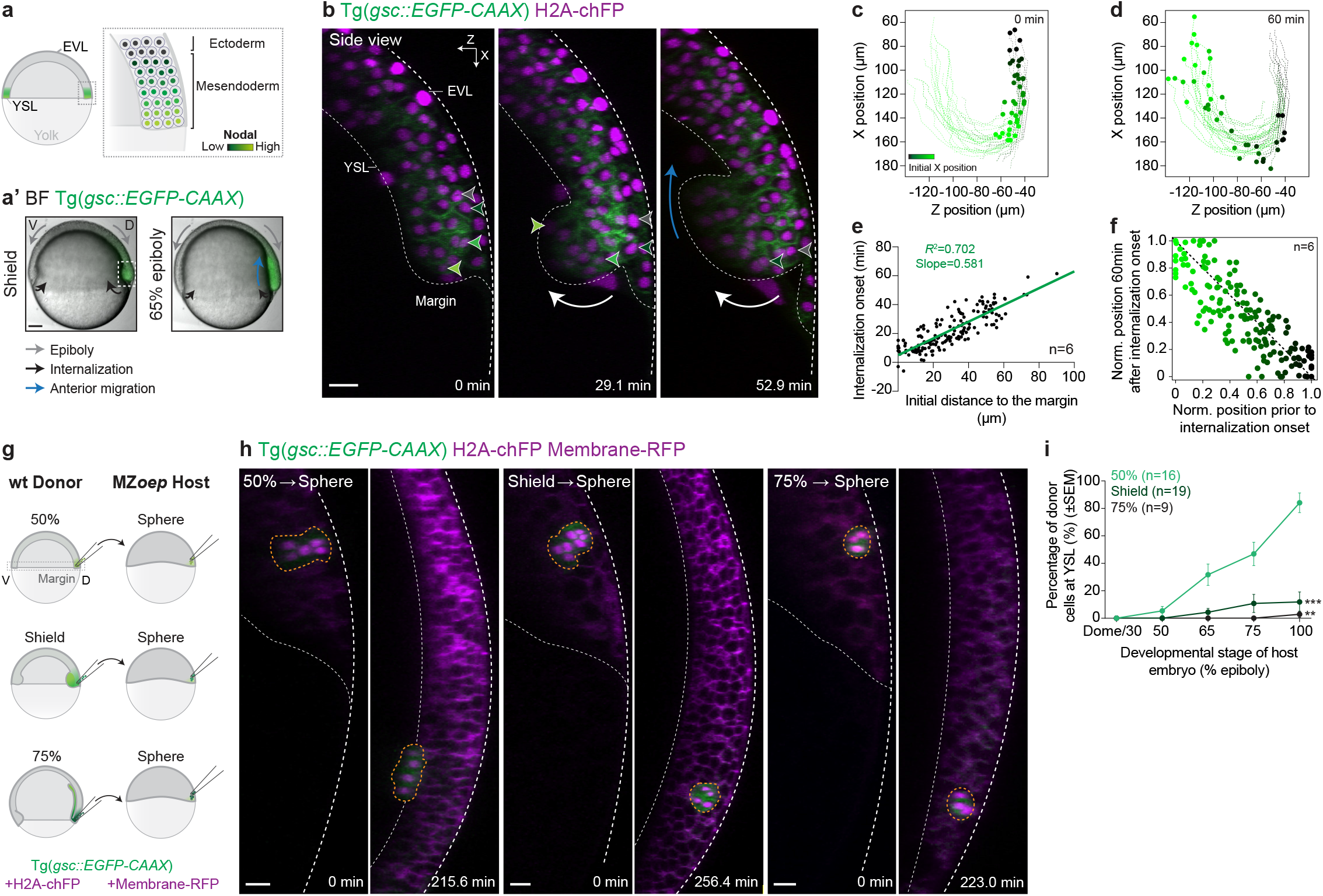
Internalization capacity of mesendoderm cells rapidly decays during gastrulation. **(a, a’)** Schematic representation of mesendoderm patterning (a) and morphogenesis (a’) during zebrafish gastrulation. Sagittal section indicating the Nodal signalling gradient (colour-coded in green) and associated embryo patterning at the onset of gastrulation (a). Bright-field single-plane images of an exemplary embryo expressing gsc::EGFP-CAAX, to mark the axial mesendoderm, soon after the onset of tissue internalization (shield stage) and during subsequent anterior migration (65% epiboly stage, a’). Arrows highlight morphogenetic movements during these stages of gastrulation. White dashed box indicates the dorsal side of the embryo, the region imaged in the subsequent high-resolution panels. **(b)** High-resolution confocal images of an embryo during mesendoderm internalization. Axial mesendoderm cells are marked by gsc::EGFP-CAAX expression (green), while H2A-chFP (magenta) marks the nuclei of all cells. Dashed lines indicate the enveloping layer (EVL) and yolk syncytial layer (YSL). White arrow highlights tissue internalization movements, while the blue arrow indicates anterior migration of internalized cells. Arrowheads indicate the relative positions of 4 exemplary cells during internalization. 0 min, internalization onset. **(c, d)** Individual tracks of internalizing mesendoderm cells shown in (b). The position of these cells is shown at the onset of internalization (c) and 60 min later (d). Colour-code corresponds to the initial distance of internalizing cells to the blastoderm margin. Plots are oriented similarly to the embryo shown in (b), with both the X and Z axes indicated in the top right corner in (b). **(e)** Onset of mesendoderm cell internalization as a function of their initial distance to the blastoderm margin in wt embryos (N=6). **(f)** Correlation between mesendoderm cell position pre- and post-internalization in wt embryos (*R*^2^=0.63, N=6; see Methods and Supplementary Note for details). Dashed line indicates perfect conservation of the relative position of cells during internalization (*R*^2^=1). Colour-code as in (c, d). **(g)** Schematic representation of the heterochronic mesendoderm cell transplants shown in (h). The dashed box highlights the blastoderm margin, the site of mesendoderm internalization *in vivo*. Note that all donor cells were both collected and then transplanted into the dorsal region of the blastoderm margin, marked by gsc::EGFP-CAAX expression. **(h)** High-resolution confocal images of exemplary wt mesendoderm donor cells, collected from the blastoderm margin of 50% epiboly (left panels), shield (middle panels) and 75% epiboly (right panels) stage embryos, transplanted into MZ*oep* hosts. Donor cells are marked by gsc::EGFP-CAAX (green) and H2A-chFP expression (magenta, nuclei), while host embryos express low levels of gsc::EGFP-CAAX (green) and Membrane-RFP (magenta). For each transplant, the first acquired time point after transplantation (0 min) and the time point when host embryos reached 100% epiboly stage are shown. Dashed white lines as in (b); yellow dashed lines highlight the donor cell transplants. **(i)** Percentage of donor mesendoderm cells, collected from 50% epiboly (N=9), shield (N=12) or 75% epiboly (N=8) stage embryos, located at the YSL as a function of the host embryo developmental stage. Kruskal-Wallis test. ***P=0.0002, **P=0.0014 (i). Lateral view (a), Dorsal view (cross-section: (b, h)). Scale bars: 100 µm (a), 20 µm (b, h).

Mesendoderm internalization is generally thought to be a cell-autonomous process, with mesendoderm cells being able to actively ingress as individuals^19,22–24^. From a biophysical perspective, differential tissue surface tension, based on differences in cell-cell adhesion and cortical tension, was originally proposed to dynamically regulate mesendoderm internalization^25–29^. More recent studies, however, suggested that these differences are not sufficient to drive mesendoderm cell internalization *in vivo,* and that this involves directed active migration^30–33^. However, it is unclear how to relate such individual internalization capacity to the highly synchronized movements observed at the tissue-scale *in vivo*^24,34,35^. Theoretical studies, supported by studies on *in vitro* cell monolayers, suggested that the interplay between single cell migration forces and supra-cellular mechanical interactions can trigger changes in collective cell behaviour resembling the jamming and glass transitions previously described for passive systems^36–38^. Thus, this raises questions as to the relative contributions of single cell mechanics, compared to collective and emergent tissue properties, to mesendoderm cell internalization *in vivo*.

Here, we show that during zebrafish gastrulation, graded Nodal signalling, in addition to its classical role as a morphogen^39–44^, encodes a mechanical switch in mesendoderm cell migratory properties via a motility-driven (un)jamming transition. This results in a collective, “leader-follower”, mode of migration which, when combined with Nodal-signalling dependent preferential adhesion, ensures ordered and pattern-preserving tissue-scale internalization movements.

## Internalization capacity of mesendoderm cells decays abruptly during gastrulation

To address how mesendoderm patterning is preserved during gastrulation, we first tracked the individual movements of dorsal mesendoderm progenitors during internalization and subsequent anterior migration (Fig. 1a’ and Extended Data Fig. 1a). In line with previous work^24,34,35,45^, we observed that mesendoderm internalization is spatially restricted to the blastoderm margin (Fig. 1b-d and Supplementary Video 1). More specifically, we found that the timing of cell internalization closely correlated with its initial distance to the margin, with cells further away internalizing later than cells right at the margin (Fig. 1e, Extended Data Fig. 1b and Supplementary Video 1). This tight link between the position and timing of cell internalization thus ensures that the first mesendoderm cells to undergo internalization will also be the first internalized cells to migrate away from the margin, thereby preserving positional information between pre- and post-internalizing progenitors (*R*^2^=0.63, Fig. 1f).

An intuitive model for linking position to internalization timing would be that the dynamics of mesendoderm cell internalization are set by an intrinsic timer, akin to classical findings on the collinear temporal activation of *Hoxb* genes in ingressing mesoderm progenitors in the chick embryo^46^. To test this possibility, we designed a system of heterochronic transplantations where mesendoderm progenitors, collected from the dorsal blastoderm margin of wild type (wt) donor embryos at different stages of gastrulation, were transplanted into the superficial layers of the blastoderm of sphere stage maternal-zygotic *oep* mutant (MZ*oep*) host embryos, which lack most mesendoderm progenitors and internalization movements^34,47,48^ (Fig. 1g and Extended Data Fig. 1a). With this, the autonomous migratory capacity of the donor cells can be distinguished from collective effects arising during normal gastrulation movements. Given that all mesendoderm donor cells were collected at the time and position where they internalize *in vivo* (i.e. the blastoderm margin), according to the timer hypothesis, all cells would thus be expected to internalize with similar dynamics (irrespective of the developmental stage of the donor embryo). In striking contrast, however, we found that only mesendoderm cells collected from early donor embryos (50% epiboly stage) efficiently internalized by the end of host embryo gastrulation (Fig. 1h, i, Extended Data Fig. 1c and Supplementary Video 2). Cells collected from donor embryos at later stages of gastrulation (shield and 75% epiboly stages), however, failed to undergo internalization and, instead, remained in superficial positions within the blastoderm (Fig. 1h, i, Extended Data Fig. 1d, e and Supplementary Video 2). Importantly, this abrupt loss of internalization competence of mesendoderm progenitors appears to be autonomous, as it was independent of the initial number of transplanted cells (Extended Data Fig. 1f), the initial distance of the transplanted cells to the host embryo margin (Extended Data Fig. 1f’) or the developmental mismatch between donor and host embryos (Extended Data Fig. 1g-i). Altogether, these findings argue for a window of competence for autonomous mesendoderm cell internalization that is closed by mid-to-late shield stage, well before the completion of internalization movements at the tissue scale. This contrasts with the view that zebrafish mesendoderm progenitors internalize simply via autonomous and synchronized ingression of single cells^22–24^ and suggests that collective properties might be involved.

## A progressive decrease in mesendoderm cell protrusiveness underlies the temporal switch of internalization capacity

To elucidate the cellular and biophysical basis for the rapid loss of autonomous internalization competence, we analysed the migratory behaviour of dorsal mesendoderm cells at different stages of gastrulation. We reasoned that this rapid loss of autonomous internalization capacity could be due to a loss of cell polarization towards the yolk syncytial layer (YSL) and/or decreasing motility forces, eventually rendering mesendoderm cells unable to drive the local cell-cell rearrangements necessary for their internalization. To discriminate between these possibilities, we transplanted small clusters of F-actin labelled internalization-competent and incompetent mesendoderm progenitors from donor embryos at 50% epiboly or shield stage, respectively, into MZ*oep* hosts, allowing us to systematically analyse their cellular protrusions in 3D over time (see Methods for details). In both internalization-competent and incompetent mesendoderm progenitors, cell protrusions were preferentially directed to the YSL (Fig. 2a, b), arguing against the notion that the loss of internalization competence in later cells is due to defective polarization. In contrast, both the number and length of cell protrusions were on average lower in internalization-incompetent mesendoderm cells (Fig. 2a, c, d, for an instantaneous analysis of cell protrusiveness see Extended Data Fig. 2b, c), suggesting that their loss of internalization competence might be linked to decreasing protrusiveness.

**Figure 2.**
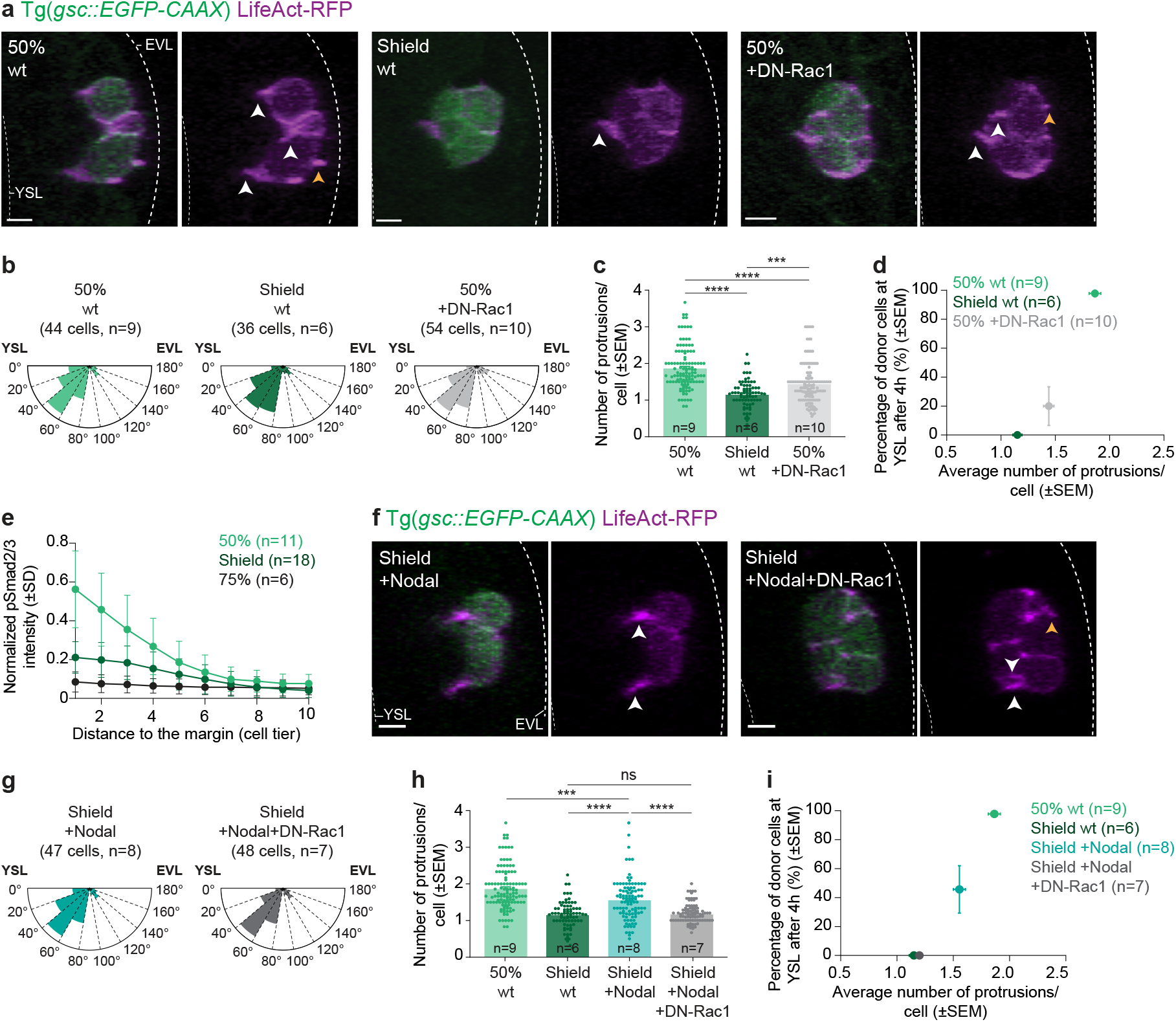
Nodal signalling regulates mesendoderm cell protrusiveness and internalization capacity. **(a)** High-resolution confocal images of exemplary wt or DN-Rac1-overexpressing donor mesendoderm cells, collected from the blastoderm margin of 50% epiboly (wt: left panels; +DN-Rac1: middle panels) or shield stage (wt: right panels) embryos and transplanted into MZ*oep* mutant host embryos. Donor cells are marked by gsc::EGFP-CAAX (green) and LifeAct-RFP expression (magenta), while host embryos express only low levels of gsc::EGFP-CAAX (green). Dashed white lines indicate the EVL and YSL. White and yellow arrowheads indicate cell protrusions oriented towards the YSL or EVL, respectively. **(b)** Rose plot of cell protrusion orientation in wt or DN-Rac1-overexpressing mesendoderm donor cells, collected from the blastoderm margin of 50% epiboly (wt: N=7; +DN-Rac1: N=10) or shield stage (wt: N=4) embryos, after transplantation into MZ*oep* hosts. **(c)** Average number of protrusions formed per wt or DN-Rac1-overexpressing mesendoderm donor cells, collected from the blastoderm margin of 50% epiboly (wt: n=44 cells, N=7; +DN-Rac1: n=54 cells, N=10) or shield stage (wt: 36 cells, N=4) embryos, after transplantation into MZ*oep* hosts. Each dot in the graph corresponds to the number of protrusions per donor cell in a single transplanted cluster at a given time point (each donor cell cluster was analysed every 5min for 60min, see Methods for details). The instantaneous number of protrusions per cell is shown for individual transplants in Extended Data Fig. 2b, c. **(d)** Percentage of wt or DN-Rac1-overexpressing mesendoderm donor cells, collected from the blastoderm margin of 50% epiboly (wt: 44 cells, N=7; +DN-Rac1: 54 cells, N=10) or shield stage (wt: 36 cells, N=4) embryos, which have arrived at the YSL of MZ*oep* hosts 4 h after the start of image acquisition, as a function of the average number of protrusions formed by these cells. **(e)** Normalized intensity of nuclear pSmad2/3 as a function of their distance to the blastoderm margin, expressed as cell tiers, in wt embryos at 50% epiboly (N=5), shield (N=6) and 75% epiboly (N=3) stage (see Extended Data Fig. 2h for exemplary images and the Methods for details). **(f)** High-resolution confocal images of exemplary mesendoderm donor cells overexpressing Nodal, alone (left panels) or in combination with DN-Rac1 (right panels), collected from the blastoderm margin of shield stage embryos and transplanted into MZ*oep* hosts. Dashed white lines and arrowheads as in (a). **(g)** Rose plot of cell protrusion orientation in mesendoderm donor cells overexpressing Nodal, alone or in combination with DN-Rac1, collected from the blastoderm margin of shield stage embryos and transplanted into MZ*oep* hosts (+Nodal: N=5; +Nodal+DN-Rac1: N=7). **(h)** Average number of protrusions formed per wt or Nodal-overexpressing mesendoderm donor cells, collected from the blastoderm margin of control and DN-Rac1 expressing embryos at 50% epiboly (wt: 44 cells, N=7) or shield stage (wt: 36 cells, N=4; +Nodal: 47 cells, N=5; +Nodal+DN-Rac1: 48 cells, N=7), and transplantated into MZ*oep* hosts. The wt data are also shown in (c). Each dot in the graph corresponds to the number of protrusions per donor cell in a single transplanted cluster at a given time point (each donor cell cluster was analysed every 5min for 60min, see Methods for details). **(i)** Percentage of wt and mesendoderm donor cells overexpressing Nodal, alone or in combination with DN-Rac1, collected from the blastoderm margin of 50% epiboly (wt: 44 cells, N=7) or shield stage (wt: 36 cells, N=4; +Nodal: 47 cells, N=5; +Nodal+DN-Rac1: 48 cells, N=7) embryos, which have arrived at the YSL of MZ*oep* hosts 4 h after the start of image acquisition, as a function of the average number of protrusions formed by these cells (see the Methods for details). Kruskal-Wallis test. ****P<0.0001 (c, h), **P=0.0012 (c), NS: Not significant (c). ***P=0.0007 (h) (c, h). Dorsal view (cross-section: (a, f)). Scale bars: 10 µm (a, f).

We thus reasoned that a critical value of protrusive/motility forces might be necessary to drive the cell-cell rearrangements necessary for mesendoderm cell internalization. Interestingly, such behaviour resembles a theoretically proposed phenomenon termed motility-driven (un)jamming^49^. To test whether this model can be applied to mesendoderm cell internalization, we first compared the mean squared displacement (MSD) of mesendoderm progenitors transplanted from 50% epiboly or shield stage donor embryos into MZ*oep* host embryos. In line with the notion that internalization-competent and incompetent mesendoderm progenitors are above or below the critical motility forces, respectively, we found that cells collected from shield stage donor embryos, in contrast to early cells (50% epiboly), displayed strongly caged motion, with limited rearrangements even over time scales of several hours (Extended Data Fig. 2d). To more directly challenge this hypothesis, we then expressed low levels of a dominant-negative version of Rac1 (DN-Rac1), a GTPase implicated in regulating mesendoderm motility^31,32,50^, in early mesendoderm progenitors to reduce protrusion formation to levels comparable to those found at later stages of gastrulation. DN-Rac1-expressing mesendoderm cells, collected from 50% epiboly stage donor embryos, displayed both reduced average number and length, but not orientation, of cell protrusions (Fig. 2a-c and Extended Data Fig. 2a). In contrast to this mild reduction in cell protrusiveness, their internalization capacity when transplanted into MZ*oep* host embryos was drastically diminished (Fig. 2d and Extended Data Fig. 2e-g’). This supports that the degree of protrusion formation is critical for the internalization competence of mesendoderm cells, and further suggests that reducing the motility forces of mesendoderm progenitors below a critical value is sufficient to jam these cells.

## Nodal signalling regulates mesendoderm cell protrusiveness and internalization capacity via a motility-driven (un)jamming transition

Next, we asked how this temporal change in mesendoderm protrusiveness, and thus internalization competence, is regulated on a molecular level. Since Nodal signalling was previously implicated in regulating mesendoderm internalization^22,23,33,47^, we reasoned that changes in Nodal signalling levels might be involved. To address this, we examined the nuclear accumulation of phosphorylated Smad2/3 complexes (pSmad2/3), a proxy for Nodal signalling activation^13–17^, in mesendoderm progenitors at different stages of gastrulation. In line with previous observations^51–55^, pSmad2/3 nuclear accumulation was highest at the blastoderm margin of embryos at the onset of gastrulation (50% epiboly), forming a steep gradient peaking at the margin and decaying towards the animal pole of the gastrula (Fig. 2e and Extended Data Fig. 2h). Notably, the peak levels of nuclear pSmad2/3 decreased in pre-internalizing progenitors at subsequent stages of gastrulation (Fig. 2e and Extended Data Fig. 2h).

To directly test whether these temporal changes in morphogen signalling might function as a control parameter tuning mesendoderm protrusiveness, and thus internalization competence, we uniformly increased Nodal ligand expression in donor embryos (Extended Data Fig. 3a, b)^28^, and transplanted these Nodal-overexpressing progenitors from shield stage donors into MZ*oep* host embryos. We found that these cells not only exhibited homogeneously higher Nodal signalling than stage-matched wt cells (Extended Data Fig. 3c-f), but also increased average number and length of mesendoderm cell protrusions (Fig. 2f-h, Extended Data Fig. 3l). Importantly, this moderate increase in mesendoderm cell protrusiveness was accompanied by a strong increase in the fraction of donor cells able to autonomously internalize, consistent with the idea that a critical value of cell protrusiveness is necessary for cell internalization (Fig. 2i and Extended Data Fig. 3g-k). To further challenge the functional relationship between Nodal signalling, cell protrusiveness and internalization competence, we co-expressed low amounts of DN-Rac1 in Nodal-overexpressing mesendoderm cells to block the enhancing effect of increased Nodal signalling on protrusion formation. Both cell protrusion formation and internalization capacity in these cells dropped to levels similar to uninduced stage-matched wt cells (Fig. 2f-i and Extended Data Fig. 3l), supporting that Nodal signalling determines the internalization competence of mesendoderm cells by regulating their protrusiveness.

To quantitatively understand the non-linear relationship between mesendoderm cell protrusiveness and internalization competence, we examined a toy-model of motility-driven (un)jamming^49^, where mesendoderm cell clusters experience a bistable energy landscape (i.e. must overcome an energy barrier to drive cellular rearrangements within the host tissue) and can move inwards, due to directed motility forces (Fig. 3a). In this simple model, mesendoderm cell clusters can locally unjam, and thus are internalization-competent, if their directed motility force overcomes a critical value (Extended Data Fig. 4a-a’’). When taking into account the measured temporal stochasticity in cell protrusiveness *in vivo* (±20% of normalized variance in protrusion number/cell, Extended Data Fig. 4b-d) via an Ornstein-Uhlenbeck equation for cell motility forces (see Methods and Supplementary Note for details), the model predicted a phase diagram with three distinct regions: for low or high average protrusive/motility forces, simulated mesendoderm cells have respectively close to 0% or 100% probability of undergoing internalization, whereas in a narrow region around the critical point, internalization outcomes are intermediate and characterized by high variability (Extended Data Fig. 4a’, e-g). This variability around the critical point emerges since clusters with average migration forces below, yet close to, the threshold can still transiently exert larger forces, allowing them to internalize (Extended Data Fig. 4c, d for examples of typical simulated trajectories; for comparison, the instantaneous number of protrusions per cell is plotted in Extended Data Fig. 2b, c). Importantly, these quantitative predictions closely matched the data across all experimental conditions, both for the average (Fig. 3b) and individual (Extended Data Fig. 4e) internalization outcomes as a function of mesendoderm donor cell protrusiveness (see Supplementary Note for details). This model also captured well, both qualitatively and quantitatively, the previously measured MSD of internalization-competent versus incompetent cells (Fig. 3c, see Supplementary Note for details), further supporting the notion that changes in mesendoderm cell protrusiveness are sufficient to account for the observed sharp changes in internalization outcomes and their associated dynamics.

**Figure 3.**
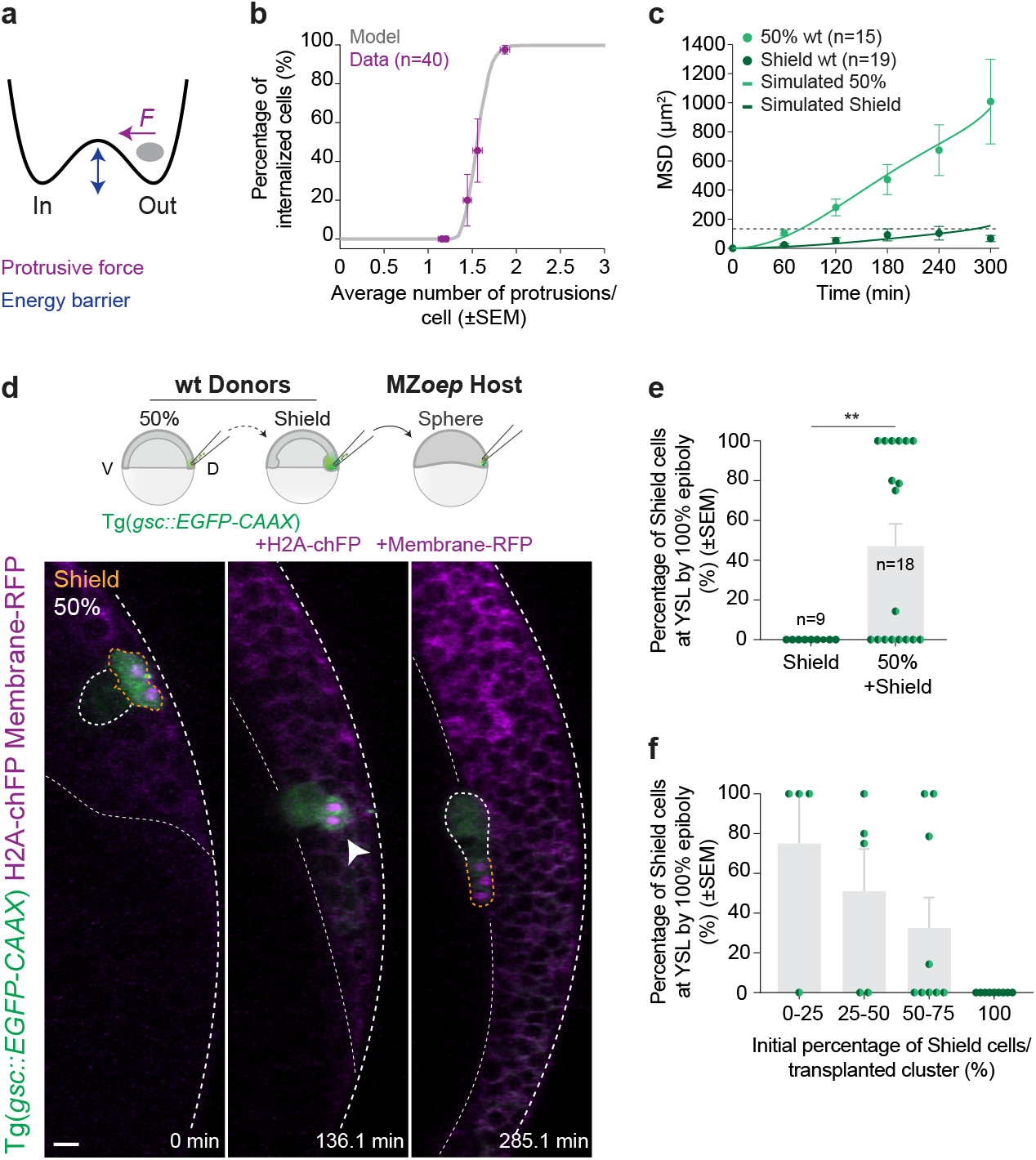
Early mesendoderm cells are above the motility-driven unjamming threshold and can act as leaders pulling on late follower cells. **(a)** Schematic representation of a toy-model for motility-driven (un)jamming (see Supplementary Note for details). **(b)** Percentage of internalized donor cells as a function of the average number of protrusions formed per cell across all experimental conditions and replicates (purple) and predicted by the model (grey, see Supplementary Note for details). The experimental data shown in the graph correspond to control or DN-Rac1-overexpressing donor mesendoderm cells, collected from the blastoderm margin of wt and Nodal-overexpressing embryos at 50% epiboly (wt: 44 cells, N=7; +DN-Rac1: 54 cells, N=10) or shield stage (wt: 36 cells, N=4; +Nodal: 47 cells, N=5; +Nodal+DN-Rac1: 48 cells, N=7), also shown in Fig. 2d, i and as individual transplants in Extended Data Fig. 4e. **(c)** Mean squared relative displacement of donor mesendoderm cells, collected from the blastoderm margin of 50% (N=9) or shield stage (N=12) embryos, or predicted by the model based on their experimentally measured average protrusiveness (n=2000). The dashed line corresponds to the average cell size at 300 min (see Methods and Supplementary Note for details). The corresponding percentage of donor cells which have arrived at the YSL as a function of host developmental stage for these transplants is shown in Fig. 1i, and the average number of cell protrusions formed per cell is shown in Fig. 2c. **(d)** High-resolution confocal images of exemplary co-transplanted mesendoderm donor cells, collected from the blastoderm margin of 50% epiboly and shield stage embryos and transplanted into a MZ*oep* mutant host embryo. Schematic representation of the co-transplantation experiments on top. All donor cells express *gsc*::EGFP-CAAX (green) and can be distinguished by H2A-chFP expression (magenta, nuclei). Host embryos express low levels of *gsc*::EGFP-CAAX (green) and Membrane-RFP (magenta). Dashed white lines indicate EVL and YSL. White and yellow dashed lines outline donor cells collected from 50% epiboly and shield stage embryos, respectively. White arrowhead points at co-transplanted donor cells forming a cohesive heterotypic cluster. 0 min, start of acquisition after transplantation. **(e)** Percentage of mesendoderm donor cells, collected from the blastoderm margin of shield stage embryos, which have arrived at the YSL of MZ*oep* hosts by the end of epiboly, when transplanted alone or together with early donor cells, collected from 50% epiboly stage embryos (N=9). **(f)** Percentage of mesendoderm donor cells collected from the blastoderm margin of shield stage embryos, which have arrived at the YSL by the end of MZ*oep* hosts epiboly, as a function of the initial composition of the co-transplanted clusters (n=18, N=9). Note that for the analyses in (e, f), only heterotypic clusters, which remained cohesive until the host embryos reached 100% epiboly, were included (Extended Data Fig. 4j, see Methods for details). Mann-Whitney test. **P=0.0084 (e). Dorsal view (cross-section: (d)). Scale bar: 20 µm (d).

Finally, we asked whether mechanisms other than the regulation of cell protrusiveness, such as cell-cell adhesion and cortical contractility, previously implicated in controlling supracellular motility transitions in *in vitro* cell monolayers^49,56,57^, might affect the competence for autonomous mesendoderm cell internalization. Measuring donor cell cluster compaction across the different conditions described, as a readout for the balance of cell-cell adhesion and contractility^58^, revealed no systematic correlation with internalization competence (Extended Data Fig. 4h, i), suggesting that Nodal signalling-dependent control of protrusiveness constitutes the central mechanism controlling the cell-autonomous capacity of mesendoderm progenitors to autonomously internalize.

## Early internalization-competent mesendoderm progenitors can act as leaders to mechanically pull late internalization-incompetent follower cells inwards

Our data so far suggest that only mesendoderm progenitors above a threshold of Nodal signalling, and thus protrusive forces, can locally unjam and autonomously internalize. Given that Nodal forms a signalling gradient peaking at the margin^51–55^ and decaying in time (Fig. 2e and Extended Data Fig. 2h), this would suggest that only a small population of cells, spatially and temporally restricted to the margin at early gastrulation stages, can drive the unjamming/cell rearrangements required to initiate internalization movements. But how would then later cells undergo internalization? One possibility is that mesendoderm cells undergo collective migration at the tissue-scale, where highly protrusive early internalization-competent cells would act as “leaders” that can mechanically pull internalization-incompetent “follower” mesendoderm cells inwards.

To test this notion experimentally, we co-transplanted marginal mesendoderm cells from 50% epiboly (internalization-competent cells, Fig. 1h, i and Supplementary Video 2) and shield stage donor embryos (internalization-incompetent cells, Fig. 1h, i and Supplementary Video 2) into MZ*oep* host embryos (Fig. 3d, top panel). Remarkably, we found that early and late cells migrated as a heterotypic cell cluster towards the inside of the blastoderm, with early cells preferentially positioned at the leading edge of the cluster, thus reaching the yolk cell first and late cells following behind (Fig. 3d, e, Extended Data Fig. 4j and Supplementary Video 3). As expected from a leader-follower organization, the likelihood of the heterotypic cluster to successfully internalize depended on their proportion of early-to-late cells, with clusters mainly composed of early cells being more likely to internalize, and clusters mainly composed of late cells remaining at the blastoderm surface (Fig. 3f). Moreover, we found that when heterotypic clusters split at the boundary between early and late cells, early cells still internalized, while late cells remained at the blastoderm surface (Extended Data Fig. 4j, k), consistent with late cells being mechanically pulled by early cells to overcome local energy barriers and collectively migrate inwards.

## Collective mesendoderm migration initiates highly ordered and pattern-preserving internalization movements *in vivo*

To determine the physiological relevance of our observations in cell transplantation assays for understanding endogenous mesendoderm internalization, we first analysed cell protrusion formation in internalizing dorsal mesendoderm cells, mosaically labelled for F-actin, during *in vivo* gastrulation. Consistent with our previous observations from the transplantation experiments and our temporal analysis of Nodal signalling *in vivo* (Fig. 2a-i, Extended Data 2a-h and 3c-l), we found a higher average number and length of protrusions in the marginal mesendoderm cells internalizing early compared to later stages of gastrulation (Extended Data Fig. 5a-f).

Next, we asked whether the parameters derived from our transplantation experiments and *in vivo* Nodal signalling gradient measurements could reproduce the large-scale internalization movements *in vivo*. To mimic mesendoderm cell internalization in a cross-section of the blastoderm margin, we employed a minimal numerical framework, previously used to study active (un)jamming in cell monolayers^59–61^, and simulated cells as motile adhesive particles in two-dimensions. In this minimal model, cells are subject to the main morphogenetic forces acting at the blastoderm margin during gastrulation: forces driving epiboly movements (F_Epiboly_), internalization movements (F_Internalization_), and anterior-directed migration of internalized mesendoderm progenitors at the YSL boundary (F_Anterior migration_) (Fig. 4a, Extended Data Fig. 5g-q and see Supplementary Note for details).

**Figure 4.**
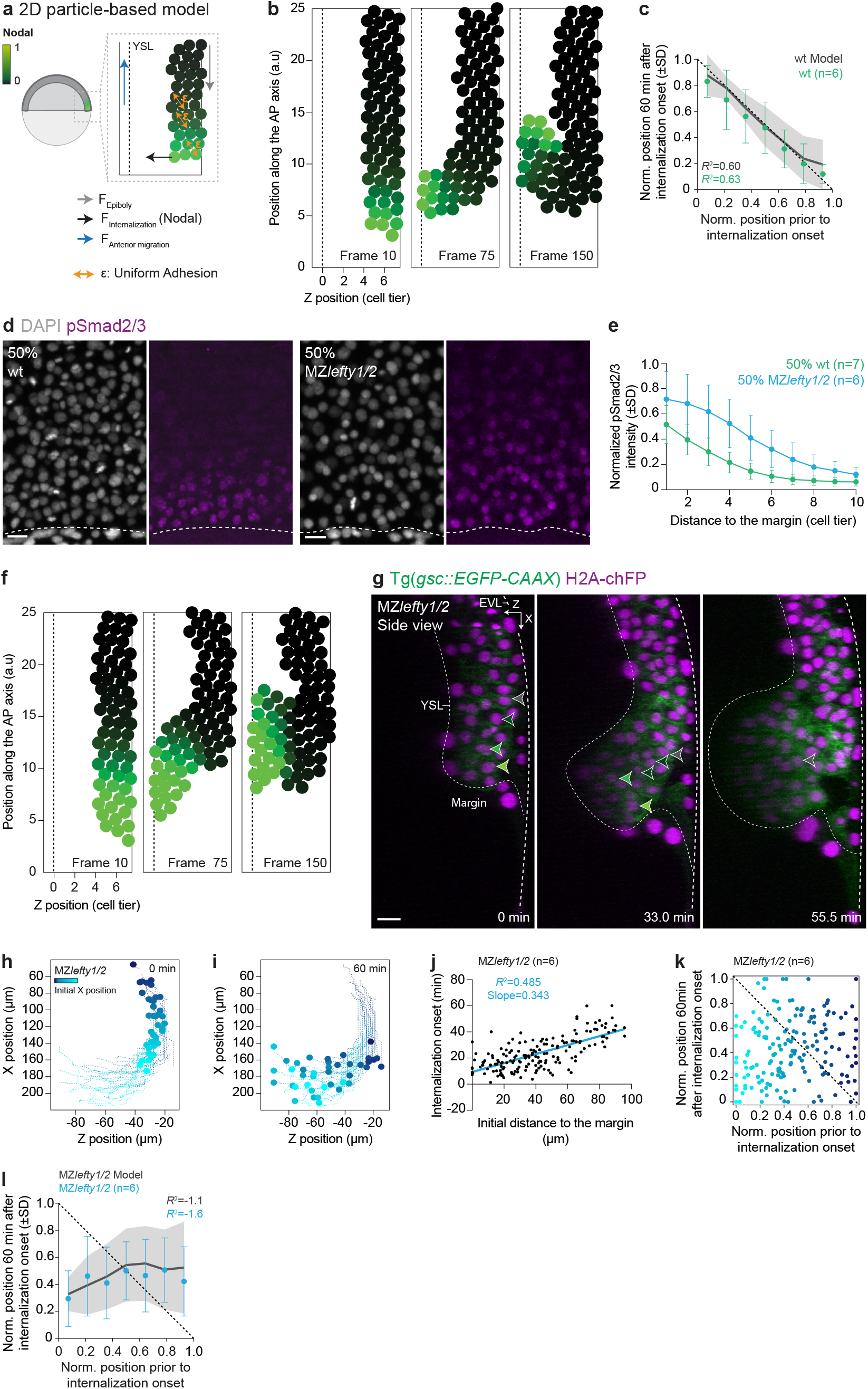
A spatiotemporal pattern of leader-to-follower cells, encoded by the Nodal signalling gradient, initiates robust mesendoderm internalization. **(a)** Schematic representation of the 2D-particle based model used to simulate mesendoderm internalization (see Supplementary Note for details). Particles are colour-coded for Nodal signalling activity. Dashed line indicates YSL. **(b)** Numerical simulations of mesendoderm internalization, based on the experimentally-measured Nodal signalling gradient at the onset of gastrulation in wt embryos (see Supplementary Note for details). Colour-code and dashed line as in (a). **(c)** Correlation between mesendoderm cell position pre-and post-internalization in wt simulations (*R*^2^=0.60, N=20 simulations) and wt embryos (*R*^2^=0.63, N=6; see Methods and Supplementary Note for details). Dashed line indicates perfect conservation of the relative position of mesendoderm cells during internalization (*R*^2^=1). The data shown for wt embryos are also plotted, as individual cells in Fig. 1f. **(d)** High-resolution confocal images of exemplary wt and MZ*lefty1/2* embryos stained for DAPI (grey, nuclei) and pSmad2/3 (magenta) at 50% epiboly stage. Dashed lines indicate deep cell margin. **(e)** Normalized intensity of nuclear pSmad2/3 as a function of their distance to the blastoderm margin, expressed as cell tiers, in wt and MZ*lefty1/2* embryos at 50% epiboly (N=3; see Methods for details). **(f)** Numerical simulations of mesendoderm internalization, based on the experimentally-measured Nodal signalling gradient at the onset of gastrulation in MZ*lefty1/2* embryos (see Methods for details). Colour-code and dashed line as in (a). **(g)** High-resolution confocal images of an exemplary MZ*lefty1/2* embryo during tissue internalization. Axial mesendoderm cells are marked by gsc::EGFP-CAAX expression (green), while all cell nuclei are marked by H2A-chFP (magenta). Dashed white lines indicate the EVL and YSL. Arrowheads indicate the relative positions of 4 exemplary cells during internalization. Note that the cell highlighted with the dark green arrow undergoes division and the position of both its daughters during internalization is subsequently indicated. 0 min, internalization onset. **(h, i)** Individual tracks of mesendoderm internalizing cells shown in (g). The position of these cells is indicated at the onset of internalization (h) and 60 min later (i). Plots are oriented similarly to the embryo shown in (g), with both the X and Z axes indicated in the top right corner in (g). Colour-code corresponds to the initial distance of internalizing cells to the blastoderm margin. **(j)** Onset of mesendoderm cell internalization as a function of their initial distance to the blastoderm margin in MZ*lefty1/2* embryos (N=6). **(k)** Correlation between cell position preand post-internalization in MZ*lefty1/2* embryos (N=6, see Methods and Supplementary Note for details). Dashed line as in (c) and colour-code as in (h, i). **(l)** Correlation between mesendoderm cell position pre- and post-internalization in MZ*lefty1/2* simulations (*R*^2^=-1.1, N=20 simulations) and MZ*lefty1/2* embryos (*R*^2^=-1.6, N=6; see Methods and Supplementary Note for details). Dashed line as in (c). The data shown for MZ*lefty1/2* embryos are also plotted as individual cells in (k). Dorsal view (top view: (d); cross-section: (g)). Scale bars: 20 µm (d, g).

We first simulated a scenario where all mesendoderm cells have equally high internalization forces. In this case, internalization was not restricted to the margin, as all mesendoderm cells were able to locally drive re-arrangement/unjamming and near-simultaneously internalize, thus resulting in a complete loss of positional order (Extended Data Fig. 5j). Conversely, lowering all internalization forces below a threshold impaired mesendoderm internalization, as cells were unable to move and displace relative to one another (Extended Data Fig. 5k). Importantly, when we inputted the experimentally measured Nodal signalling gradient at the onset of gastrulation, and assumed a linear relationship between Nodal signalling and internalization forces (as supported by the cell transplantation assays and analysis of Nodal signalling dynamics *in vivo*, see Fig. 2a-i, Extended Data 2a, h, 3c-l and 5a-g and Supplementary Note for details), we found that only the marginal-most cells had sufficiently high forces to locally unjam. This restricted tissue-scale mesendoderm internalization to the margin, with internalization-competent leader cells rapidly reaching the YSL boundary and pulling the remaining cells towards the inside (Fig. 4b and Supplementary Video 4). Notably, these simulations closely reproduced the initiation of tissue internalization observed *in vivo* (Fig. 1b, 4b, Supplementary Videos 1 and 4) and quantitatively matched the degree of positional order preservation observed experimentally (respectively *R*^2^=0.60 versus *R*^2^=0.63, Fig. 4c, see Supplementary Note for further details on statistics). This further supports the notion that the Nodal signalling gradient, by subdividing the mesendoderm tissue into leader and follower cells, initiates highly ordered, pattern-preserving internalization movements at the blastoderm margin.

## Nodal signalling-encoded ratio of leader and follower cells is critical for ordered tissue internalization *in vivo*

To further challenge this model, we asked whether the shape of the Nodal signalling gradient at the blastoderm margin is critical for ordered mesendoderm internalization. To this end, we examined maternal-zygotic mutants of the Nodal inhibitor *lefty1/2* (MZ*lefty1/2*), which exhibit increased Nodal signalling and mesendoderm specification^62^. Consistent with previous findings^62^, our quantitative analysis indicated that MZ*lefty1/2* embryos display higher peak levels of Nodal signalling and an expanded gradient within the blastoderm margin (Fig. 4d, e and Extended Data Fig. 6a, b). Given our previous findings on the relationship between Nodal signalling, mesendoderm cell protrusiveness, we reasoned that this expanded Nodal gradient should increase the number of leader cells above the critical motility force required for local unjamming and autonomous internalization. When using the experimentally-measured Nodal signalling gradient in MZ*lefty1/2* embryos as an input parameter for F_Internalization_ in our particle-based model (all remaining parameters were kept unchanged; see Supplementary Note for details), simulations predicted a highly disorganized internalization process where positional order was lost (Fig. 4f and Supplementary Video 4). This was due to an increased number of leader cells simultaneously internalizing at multiple locations, in contrast with the wild type scenario of highly restricted (un)jamming at the blastoderm margin (*R*^2^=-1.1, Fig. 4f, l, Supplementary Video 4 and see Supplementary Note for details). To test these predictions experimentally, we tracked the individual movements of mesendoderm progenitors in MZ*lefty1/2* embryos at the onset of internalization and found a clearly more disorganized internalization process than in wt embryos (respectively *R*^2^=-1.6 versus *R*^2^=0.63, Fig. 4g-k, Extended Data Fig. 6c, Supplementary Video 5 and Fig. 1b-f, Extended Data Fig. 1b, Supplementary Video 1), matching our model predictions both qualitatively and quantitatively (Fig. 4f, g, l, Supplementary Videos 4 and 5).

Finally, in line with the assumption that an expanded Nodal signalling gradient increases the number of leader cells, we found that mesendoderm cells collected from MZ*lefty1/2* embryos displayed a prolonged competence to unjam and autonomously internalize upon transplantation into MZ*oep* hosts, when compared to stage-matched wt cells (Extended Data Fig. 6d-f’, l). Moreover, mesendoderm cells internalizing at the blastoderm margin on MZ*lefty1/2* embryos, mosaically labelled for F-actin, displayed a sustained increase in the average number and length of protrusions compared to wt cells (Extended Data Fig. 6h-k and 5a-f), consistent with the notion of Nodal signalling controlling the window of autonomous mesendoderm cell internalization capacity by regulating cell protrusiveness. In contrast, wt and MZ*lefty1/2* mesendoderm cells formed cell clusters of similar compaction upon transplantation (Extended Data Fig. 6g), suggesting that changes in cell-cell adhesion and cortical contractility do not play a major role in regulating internalization competence. Collectively, these findings argue that the shape of the Nodal signalling gradient is critical to set the ratio of highly protrusive leader to less protrusive follower cells, an essential feature for triggering collective and orderly mesendoderm internalization movements at the blastoderm margin.

## Nodal signalling-mediated preferential adhesion is required for preserving positional information during later stages of mesendoderm internalization

For collective and ordered mesendoderm internalization to occur at the tissue-scale, leader cells need to establish and maintain strong cell-cell contacts with their immediate followers in order to transmit pulling forces throughout internalization and subsequent anterior migration. While, at the start of mesendoderm internalization, this is easily achieved as cells are mainly in contact with their immediate neighbours, once internalization movements are underway, internalized mesendoderm cells will also come into contact with not-yet internalized progenitors over the newly formed tissue boundary. To probe whether and how positional information is maintained during these later stages of gastrulation, we extended our tracking analysis of mesendoderm internalization and anterior migration both in our particle-based simulations and *in vivo*. Interestingly, we found that positional order within the internalized mesendoderm during later stages of tissue internalization and anterior migration was increasingly lost in the simulations, but still maintained *in vivo* (respectively *R*^2^=0.33 versus *R*^2^=0.83; Fig. 5a, b, Supplementary Video 4 and see Supplementary Note for details).

**Figure 5.**
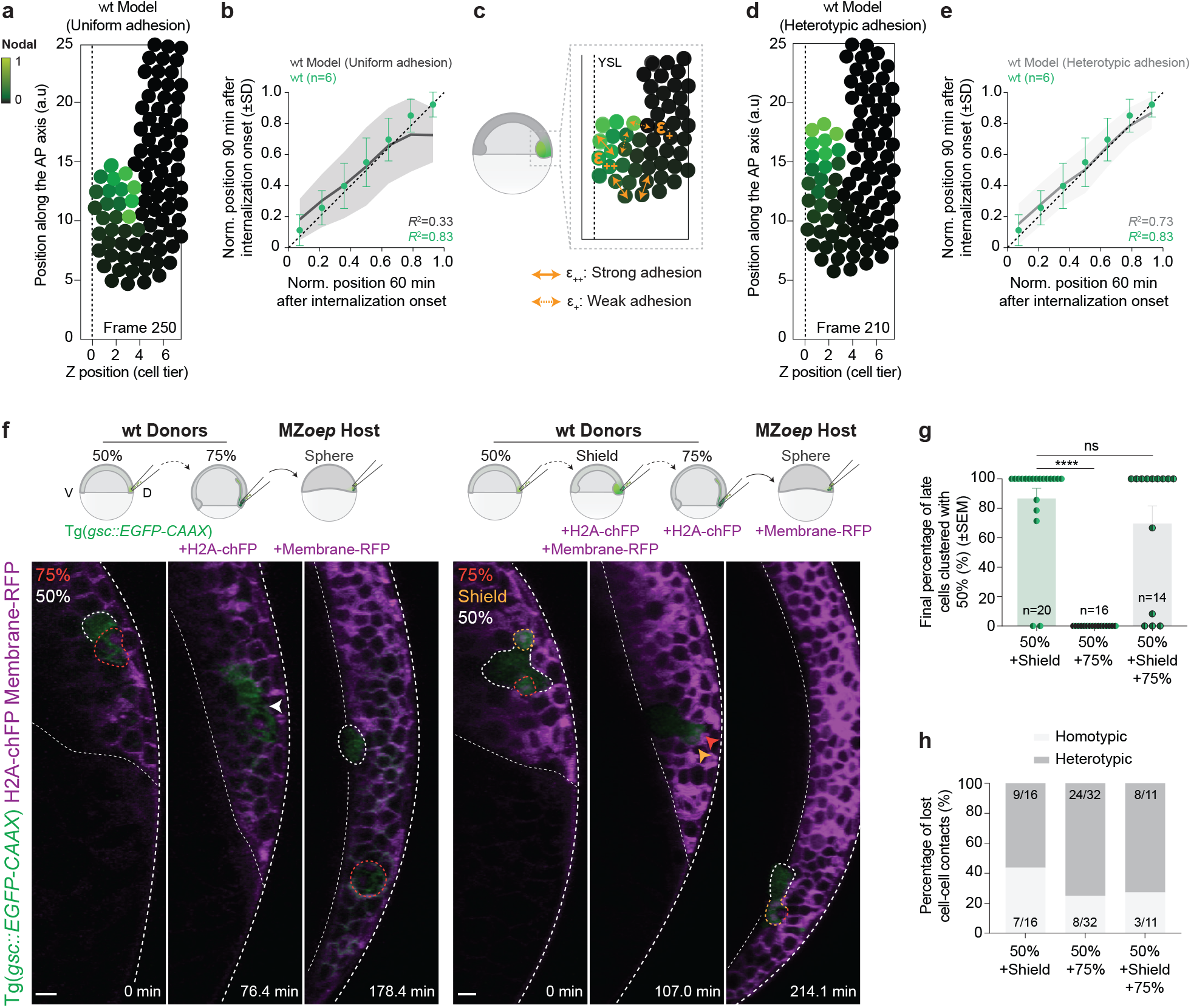
Leader/follower cohesion contributes for orderly mesendoderm internalization and migration. **(a)** Numerical simulation of mesendoderm internalization, based on the experimentally-measured Nodal signalling gradient at the onset of gastrulation in wt embryos. This corresponds to a later time point of the simulations shown in Fig. 4b (see Supplementary Note for details). Particles are colour-coded for Nodal signalling activity. Dashed line indicates the YSL. **(b)** Correlation between cell position at later stages of tissue internalization in wt simulations assuming uniform adhesion (*R*^2^=0.33, N=20 simulations) and wt embryos (*R*^2^=0.83, N=6; see Methods and Supplementary Note for details). Dashed line indicates perfect conservation of the relative position of mesendoderm cells during internalization (*R*^2^=1). **(c)** Schematic representation of the 2D-particle based model used to simulate mesendoderm internalization assuming Nodal-dependent heterotypic interactions. Colour-code and dashed line as in (a). All remaining parameters were left as in Fig. 4b (see Supplementary Note for details). **(d)** Numerical simulations of mesendoderm internalization assuming Nodal signalling-dependent heterotypic, rather than uniform, interactions (see Supplementary Note for details). Colour-code and dashed line as in (a). **(e)** Correlation between cell position at later stages of tissue internalization in wt simulations assuming Nodal signalling-dependent heterotypic interactions (*R*^2^=0.73, N=20 simulations) and wt embryos (*R*^2^=0.83, N=6; see Methods and Supplementary Note for details). Dashed line as in (b). **(f)** High-resolution confocal images of exemplary co-transplanted mesendoderm donor cells, collected from the blastoderm margin of 50% and 75% epiboly stage embryos (left) or 50%, shield and 75% epiboly stage embryos (right) and transplanted into MZ*oep* mutant host embryos. All donor cells express gsc::EGFP-CAAX (green) and can be distinguished by H2A-chFP expression (magenta, nuclei; 75% cells) or H2A-chFP and Membrane-RFP co-expression (both in magenta; shield cells). Host embryos express low levels of gsc::EGFP-CAAX (green) and Membrane-RFP (magenta). Dashed white lines indicate the EVL and YSL. White, yellow and orange dashed lines outline donor cells collected from 50% epiboly, shield and 75% epiboly stage embryos, respectively. Schematic representations of the co-transplantation assays are shown in the top. Arrowheads point at co-transplanted donor cells forming cohesive heterotypic clusters. **(g)** Percentage of co-transplanted late (Shield, 75% or Shield+75%) mesendoderm donor cells that remain clustered with early cells (50%) in mixed clusters of different compositions (50%+Shield: N=10; 50%+75%: N=9; 50%+Shield+75%: N=6) until MZ*oep* hosts reached 100% epiboly stage. **(h)** Percentage of homotypic and heterotypic cell-cell contacts lost upon the final splitting of mesendoderm donor cell clusters of different compositions (50%+Shield: N=10; 50%+75%: N=9; 50%+Shield+75%: N=6) until the MZ*oep* hosts reached 100% epiboly stage (see Methods for details). Kruskal-Wallis test. ****P<0.0001, NS: Not significant (g). Dorsal view (cross-section: (f)). Scale bars: 20 µm (f).

To understand this discrepancy, we computed average velocity maps for internalized mesendoderm cells during anterior migration. We found that in the simulations, but not *in vivo*, internalized mesendoderm cells migrating away from the blastoderm margin displayed uniformly lower velocities near the forming tissue boundary than close to the YSL (Extended Data Fig. 7a-c). This velocity gradient in the simulations was due to strong adhesion between internalized and not-yet-internalized cells, resulting in extensive cell-cell rearrangements mainly along the tissue boundary and, consequently, loss of positional order within the internalized tissue (*R*^2^=0.33; Extended Data Fig. 7a, b, Fig. 5a, b and Supplementary Video 4). In contrast to this, strong velocity gradients were only detectable at the most-marginal region of blastoderm *in vivo,* thereby preserving positional order within mesendoderm also at later stages of tissue internalization and anterior migration (*R*^2^=0.83, Extended Data Fig. 7c, Fig. 5b and Supplementary Video 4).

This divergence between model predictions and experimental observations suggests that, in contrast to our original simplified model assumption of uniform cell-cell adhesion, adhesion might instead be regulated in a position-dependent manner, thereby minimizing cross-boundary effects. Given previous findings that Nodal signalling modulates cell-cell adhesion^28,32,63–65^, we computationally addressed two possible scenarios: *differential adhesion*, where the Nodal signalling gradient is translated into a gradient of absolute adhesion strength within the mesendoderm, and *heterotypic/preferential adhesion*, where Nodal signalling determines preferential adhesion amongst mesendoderm cells, with stronger cell-cell adhesion between cells located at similar positions within the Nodal gradient and, thus similar Nodal signalling activity (Extended Data Fig. 7d, e and Fig. 5c). Interestingly, simulations incorporating heterotypic/preferential adhesion, but not differential adhesion (all remaining parameters were kept unchanged, see Supplementary Note for details), robustly produced orderly cell movements matching the experimental observations not only at the onset of internalization (*R*^2^=0.67 and *R*^2^=0.15 in simulations assuming heterotypic or differential adhesion, respectively versus *R*^2^=0.63 in wt embryos), but also during later stages of tissue internalization and anterior migration (*R*^2^=0.73 and *R*^2^=0.18 in simulations assuming heterotypic or differential adhesion, respectively versus *R*^2^=0.83 in wt embryos; Fig. 5d, e, Extended Data Fig. 7e-j, Supplementary Video 4 and see Supplementary Note for details). In line with this, simulations with heterotypic/preferential adhesion also closely reproduced the experimental mesendoderm velocity maps, with nearly uniform velocities for internalized mesendoderm cells located at a distance from the margin (Extended Data Fig. 7c, k).

To experimentally test our model assumption of preferential/heterotypic adhesion amongst mesendoderm cells, we performed a set of co-transplantation experiments, where donor cells with different Nodal signalling activity were co-transplanted into MZ*oep* embryos and monitored for their ability to preserve cluster cohesiveness, a functional and physiologically-relevant readout of cell-cell adhesion strength in this context (Fig. 5f). We reasoned that, if the difference in Nodal signalling between contacting cells indeed inversely scales with their cohesion, then the integrity of a leader-follower cluster should be preserved where differences are small and lost once these differences increase (see Extended Data Fig. 7d for a schematic). In line with this, our previous co-transplantation experiments with mesendoderm progenitors from donor embryos at 50% epiboly and shield stage (comparably small differences in Nodal signalling activity, Fig. 2e and Extended Data Fig. 2h) showed that cluster integrity was well preserved over time, as evidenced by the large proportion of shield cells undergoing internalization together with 50% epiboly cells (Fig. 3d-f, Extended Data Fig. 4j, k and Supplementary Video 3). In contrast to this, when we co-transplanted mesendoderm cells from donor embryos at 50% and 75% epiboly stage (comparably large differences in Nodal activity, Fig. 2e and Extended Data Fig. 2h), cluster integrity was lost, with early (50% epiboly) cells typically internalizing and late (75% epiboly) cells remaining jammed in superficial layers of the blastoderm (Fig. 5f, g and Supplementary Video 6). Notably, these clusters preferentially split at the boundary between early and late cells (Fig. 5h and Supplementary Video 6), consistent with our assumption that adhesion is weakest at the contacts between cells with large differences in Nodal signalling, and that strong cell-cell adhesion between neighbouring cells is required for the leaders to mechanically pull, and locally unjam, their followers.

To further test the role of Nodal signalling in this process, we again co-transplanted mesendoderm cells from 50% and 75% epiboly stage donor embryos into MZ*oep* host embryos, but this time using Nodal-overexpressing 75% epiboly stage donor embryos. With this, we kept the differences in developmental time between co-transplanted cells constant, but narrowed their overall difference in Nodal signalling activity. In line with this, Nodal-overexpressing cells at 75% epiboly stage adopt an anterior axial mesendoderm fate (Extended Data Fig. 3a, b), as expected from cells exposed to high Nodal signalling^14,17,41,42,66–68^. In these co-transplantations, cluster cohesion was strongly enhanced compared to control co-transplantations of mesendoderm cells from 50% and uninduced 75% epiboly stage donor embryos (Extended Data Fig. 7l-n), supporting that the adhesion strength at heterotypic contacts is modulated by the differences in Nodal signalling activity.

As an ultimate test to our model, we co-transplanted mesendoderm cells from 50% epiboly, shield and 75% epiboly stage donor embryos in MZ*oep* host embryos (Fig. 5f see right panel). We reasoned that in such triple clusters, shield cells should be able to mediate adhesion between early (50% epiboly) and late cells (75% epiboly), mimicking the adhesion sequence of the gastrulating embryo (see Extended Data Fig. 7d for a schematic). Strikingly, we found that the proportion of triple transplants that remain cohesive by the end of host embryo epiboly was strongly increased when compared to clusters composed solely of 50% and 75% mesendoderm cells (Fig. 5f-h). Moreover, we observed that, in some cases, cohesive triple clusters completed their internalization, with early cells (50% epiboly) positioned at the leading edge, followed by shield and then 75% epiboly stage cells (Fig. 5f see right panel), as predicted for Nodal signalling determining not only a switch in internalization competence of mesendoderm cells, but also preferential adhesion amongst mesendoderm progenitors.

## Discussion

A key finding of our study is that a morphogen gradient encodes a sharp mechanical switch in migration competence *in vivo*. The observation that below a threshold of motility force, dorsal mesendoderm cells are unable to autonomously internalize, despite remaining polarized towards the YSL, is a hallmark of a motility-driven (un)jamming transition, previously explored in cell monolayers via self-propelled vertex models^49,57^. This constitutes a complementary mechanism to recent studies of (un)jamming transitions *in vitro*^56,69^ and *in vivo*^70–76^, where cell movement/uncaging is used as a proxy for tissue unjamming, and is mechanistically controlled by changes in the adhesive and contractile properties of cells^69,71–73,76^. In our study, in contrast, graded cell motility - downstream of Nodal signalling - functions as the control parameter triggering this transition, thereby determining the ability of a subset of mesendoderm cells to locally unjam and migrate towards the YSL. This morphogen-encoded (un)jamming transition not only provides insight into the effector mechanisms by which mechanochemical signalling converts a graded upstream signal into the discrete mechanical domains necessary for collective morphogenesis, but also about the functional relevance of (un)jamming transitions in development^36–38^. Indeed, we found that this transition ensures that mesendoderm internalization remains restricted to the blastoderm margin, a prerequisite for collective and ordered tissue internalization.

Collective migration, a fundamental process in a number of developmental, homeostatic and pathological settings, promotes directed migration by strengthening the response of individual cells to extrinsic guidance cues and/or driving supracellular polarization^77–82^. Our findings point at another important function of collective migration in preserving tissue patterning during complex 3D tissue movements. A critical determinant of this collective mode of tissue internalization is the ability of leader cells to effectively pull on their immediate followers, a process dependent on cell-cell adhesion. Our data suggest that mesendoderm cells originally located at a similar position within the Nodal signalling gradient before internalization, not only acquire a similar fate^39–44^, but also display preferential adhesion. Such preferential adhesion mechanism, previously proposed to be efficient at driving cell sorting^83^, is therefore also critical for collective migration. Indeed, it provides both strong cohesion along the path of internalization, by efficiently transmitting pulling forces from leaders to their designated follower cells, and minimizing interactions across the forming tissue boundary during latter stages of internalization and anterior migration. This also suggests a dual mechanical role of Nodal signalling in mesendoderm collective migration, by controlling both cell protrusion forces to regulate leader-follower proportions and cell-cell adhesion forces to ensure cohesion between the leader and follower domains.

How the biochemical signals determining embryo patterning and the mechanical forces driving morphogenesis are integrated is still a matter of intense research. The temporal sequence by which these two processes occur in development is important, particularly when tissue morphogenesis follows patterning, since this would be expected to perturb the initial pattern. This is relevant for zebrafish gastrulation, where Nodal signalling-mediated patterning occurs primarily in blastula and early gastrula stages^18^, prior to, or concomitant with the extensive 3D migratory movements associated with germ layer internalization. Our findings identify a cross-scale mechanism where a morphogen gradient ensures pattern preservation despite large-scale morphogenesis, by functioning both as a morphogen controlling cell fate specification, and a ‘mechanogen’^84^ triggering collective migration through a motility-driven (un)jamming transition. Since morphogen gradients are highly conserved in vertebrate development^1,13,20,21,85–87^, similar mechanochemical gradients might be relevant in other developmental and disease-related processes, such as wound healing, thereby constituting a general principle coordinating tissue patterning and morphogenesis.

## Supporting information

Supplementary Theory Note

## Acknowledgments

We thank Karuna Sampath, Andrea Pauli and Yohanns Bellaїche for feedback on the manuscript. We also thank the members of the Heisenberg group, in particular Alexandra Schauer and Feyza Nur Arslan, for help, technical advice and discussions, and the Bioimaging and Life Science facilities at IST Austria for continuous support. This work was supported by postdoctoral fellowships from EMBO (LTF-850-2017) and HFSP (LT000429/2018-L2) to D.P. and the European Union (European Research Council Starting Grant 851288 to E.H. and European Research Council Advanced Grant 742573 to C.-P.H.).

## Author contributions

D.P., E.H. and C.-P.H. designed the research. D.P. performed the experiments and analysed the experimental data. R.K. created the DN-Rac1 plasmid. E.H. developed the numerical simulations and analysed the experimental data. D.P. and C.-P.H. wrote the manuscript with input from all authors.

## Supplementary Figures

**Extended Data Figure 1.**
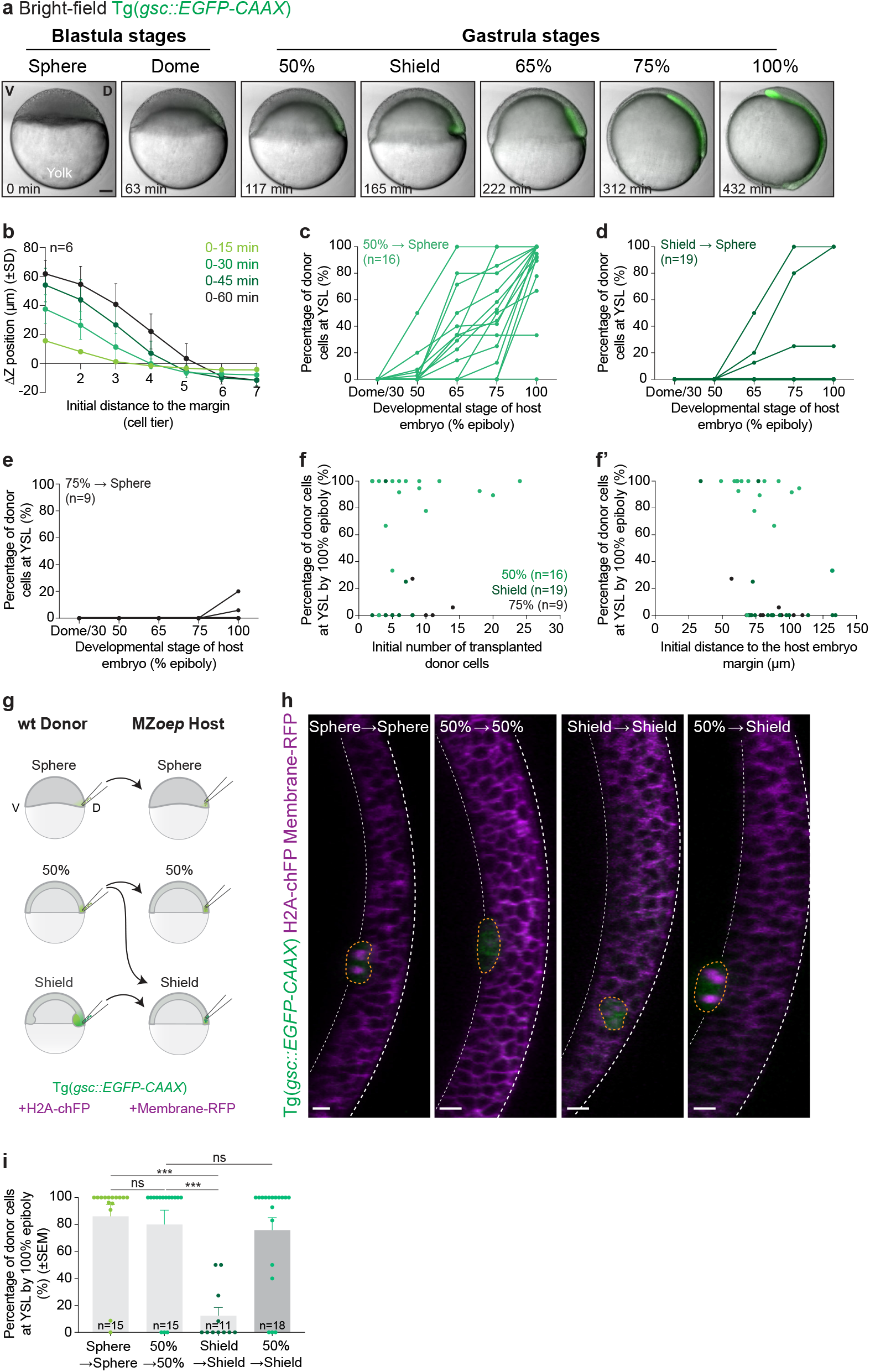
Loss of internalization capacity of mesendoderm cells during gastrulation is cell-autonomous. **(a)** Bright-field single-plane images of an exemplary embryo expressing gsc::EGFP-CAAX, marking axial mesendoderm cells, during gastrulation. **(b)** Change in Z position, a proxy for cell internalization, of mesendoderm cells as a function of their initial distance to the margin, expressed in cell tiers, in wt embryos (N=6). The colour-code corresponds to the indicated time bins. **(c-e)** Percentage of mesendoderm donor cells, collected from the blastoderm margin of 50% epiboly (c, N=9), shield (d, N=12) or 75% epiboly (e, N=8) stage embryos, which have arrived at the YSL, as a function of the host developmental stage. **(f, f’)** Percentage of donor mesendoderm cells, collected from the blastoderm margin of 50% epiboly (N=9), shield (N=12) or 75% epiboly (N=8) stage embryos, which have arrived at the YSL by the end of host embryo epiboly, as a function of the initial number of transplanted cells (f) or the initial distance of the transplanted cells to the host embryo margin (f’). **(g)** Schematic representation of the homotypic and heterochronic mesendoderm cell transplants shown in (h). **(h)** High-resolution confocal images of exemplary mesendoderm donor cells, collected from the blastoderm margin of sphere, 50% epiboly or shield stage embryos and transplanted into sphere, 50% epiboly or shield stage MZ*oep* hosts. Donor cells are marked by gsc::EGFP-CAAX (green) and H2A-chFP expression (magenta, nuclei), while host embryos express low levels of gsc::EGFP-CAAX (green) and Membrane-RFP (magenta). The stage of donor and host embryos are indicated at the top. Single time points by the end of host embryo epiboly are shown. Dashed white lines indicate EVL and YSL, while the yellow dashed line outline donor cells. **(i)** Percentage of mesendoderm donor cells, collected from the blastoderm margin of sphere, 50% epiboly or shield stage embryos, which have arrived at the YSL by the end of host embryo epiboly. Note that donor cells were transplanted into either stage-matched host embryos (donor and host embryos at sphere, N=6; 50% epiboly, N=5; donor and host embryos at shield stage, N=7) or shield stage hosts (N=7). The stage of both donor and host embryos are indicated below. Kruskal-Wallis test. NS: Not significant, \*\*\**P*=0.0004 (Sphere → Sphere vs. Shield → Shield), \*\*\**P*=0.0003 (50% → 50% vs. Shield → Shield) (i). Lateral view (a), Dorsal view (cross-section: (h)). Scale bars: 100 µm (a), 20 µm (h).

**Extended Data Figure 2.**
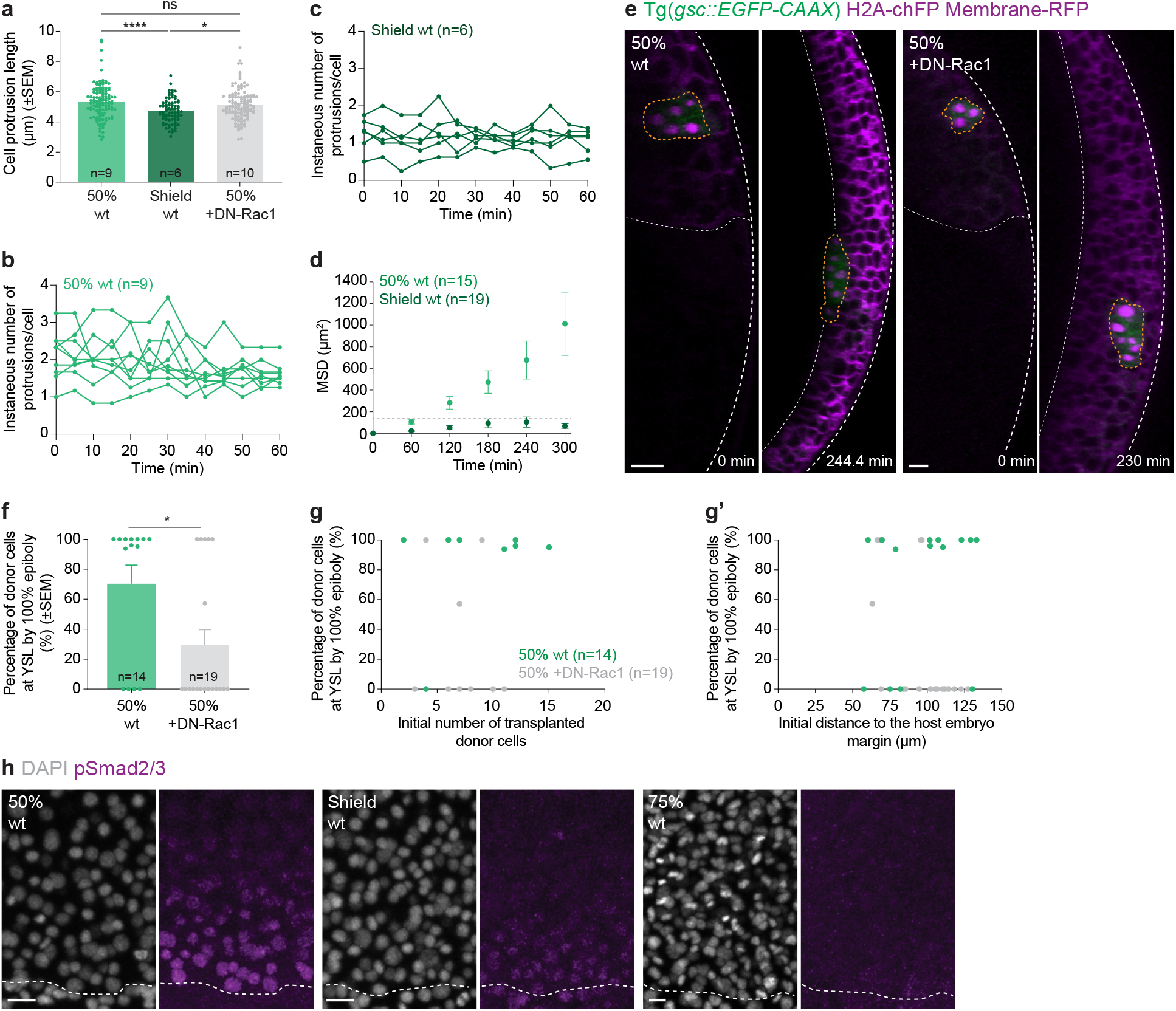
Internalization-incompetent mesendoderm donor cells exhibit caged motion. **(a)** Length of protrusions formed by mesendoderm donor cells, collected from the blastoderm margin of wt or DN-Rac1-overexpressing donor embryos at 50% epiboly (wt: 44 cells, N=7; +DN-Rac1: 54 cells, N=10) or shield stage (wt: 36 cells, N=4) and transplanted into MZ*oep* hosts. Each dot in the graph corresponds to the average length of cell protrusions in a single transplanted cluster at a given time point (each donor cell cluster was analysed every 5min for 60min, see Methods for details). **(b, c)** Instantaneous number of protrusions formed per mesendoderm donor cell, collected from the blastoderm margin of 50% (b, 44 cells, N=7) or shield stage (c, wt: 36 cells, N=4) embryos and transplanted into MZ*oep* hosts. Each curve corresponds to a single transplanted cluster (see Methods for details). **(d)** Mean squared relative displacement (MSD) of mesendoderm donor cells, collected from the blastoderm margin of 50% (N=9) or shield stage (N=12) embryos and transplanted into MZ*oep* hosts. The dashed line corresponds to the average cell size at 300 min (see Methods and Supplementary Note for details). The percentage of donor cells, which have arrived at the YSL, as a function of host developmental stage for these transplants is shown in Fig. 1i. **(e)** High-resolution confocal images of control or DN-Rac1-overexpressing mesendoderm donor cells, collected from the blastoderm margin of 50% epiboly embryos and transplanted into MZ*oep* hosts. Donor cells are marked by gsc::EGFP-CAAX (green) and H2A-chFP expression (magenta, nuclei), while host embryos express low levels of gsc::EGFP-CAAX (green) and Membrane-RFP (magenta). For each transplant, the first acquired time point and the time point when the hosts reached 100% epiboly stage are shown. Dashed white lines indicate EVL and YSL. Yellow dashed lines outline donor cells. **(f)** Percentage of control or DN-Rac1-overexpressing mesendoderm donor cells, collected from the blastoderm margin of 50% epiboly embryos, which have arrived at the YSL by the end of host embryo epiboly (N=9). **(g, g’)** Percentage of control or DN-Rac1-overexpressing mesendoderm donor cells, collected from the blastoderm margin of 50% epiboly embryos, which have arrived at the YSL by the end of host embryo epiboly, as a function of the initial number of transplanted cells (g; N=9) or the initial distance of the transplanted cells to the host embryo margin (g’; N=9). **(h)** High-resolution confocal images of exemplary wt embryos stained for DAPI (grey, nuclei) and pSmad2/3 (magenta) at 50% epiboly (left), shield (middle) and 75% epiboly (right) stage. Dashed lines indicate deep cell margin and a quantification of pSmad2/3 intensity as a function of distance to the blastoderm margin is shown in Fig. 2e. Kruskal-Wallis test. NS: not significant. ****P=<0.0001, *P=0.0102 (a). *t* test. *P=0.0159 (f). Dorsal view (cross-section: (e), top view: (h)). Scale bars: 20 µm (e, h).

**Extended Data Figure 3.**
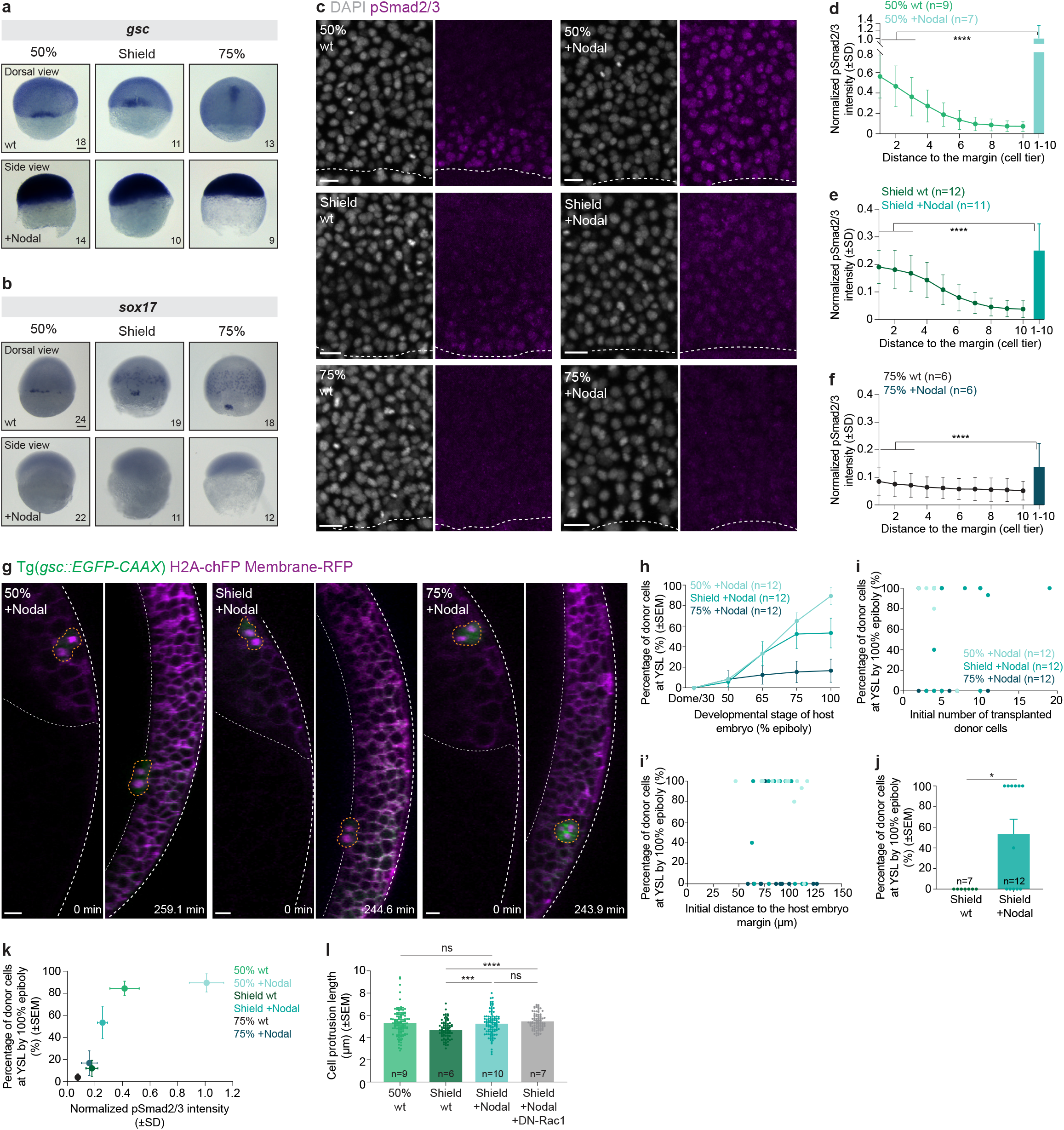
Increasing Nodal signalling in late mesendoderm cells prolongs their internalization capacity. **(a, b)** Expression of mesendoderm marker genes (*gsc* (a) and *sox17* (b)) as determined by whole mount *in situ* hybridization of wt and Nodal-overexpressing embryos at different stages of gastrulation. The number of embryos per condition is indicated in the lower right corner (N=3 (a); N=2 (b)). **(c)** High-resolution confocal images of exemplary wt and Nodal-overexpressing embryos stained for DAPI (grey, nuclei) and pSmad2/3 (magenta) at 50% epiboly (top), shield (middle) and 75% epiboly (bottom) stage. Dashed lines indicate deep cell margin. **(d-f)** Normalized intensity of nuclear pSmad2/3 as a function of their distance to the blastoderm margin, expressed as cell tiers, in wt and Nodal-overexpressing embryos at 50% epiboly (d, N=4), shield (e, N=4) and 75% epiboly (f, N=3) stage (see Methods for details). **(g)** High-resolution confocal images of exemplary Nodal-overexpressing donor cells, collected from the blastoderm margin of 50% epiboly (left), shield (middle) and 75% epiboly (right) stage embryos and transplanted into MZ*oep* mutant host embryos. Donor cells are marked by gsc::EGFP-CAAX (green) and H2A-chFP expression (magenta, nuclei), while host embryos express low levels of gsc::EGFP-CAAX (green) and Membrane-RFP (magenta). For each transplant, the first acquired time point and the time point when hosts reached 100% epiboly stage are shown. Dashed white lines indicate EVL and YSL; yellow dashed lines outline donor cell transplants. **(h)** Percentage of Nodal-overexpressing donor cells, collected from 50% epiboly (N=5), shield (N=4) or 75% epiboly (N=6) stage embryos, which have arrived at the YSL, as a function of the host embryo developmental stage. **(i, i’)** Percentage of Nodal-overexpressing donor cells, collected from the blastoderm margin of 50% epiboly (N=5), shield (N=4) or 75% epiboly (N=6) stage embryos, which have arrived at the YSL by the end of MZ*oep* host embryo epiboly, as a function of the initial number of transplanted cells (i) or the initial distance of the transplanted cells to the host embryo margin (i’). **(j)** Percentage of control and Nodal-overexpressing donor cells, collected from the blastoderm margin of shield (N=4) stage embryos, which have arrived at the YSL by the end of host embryo epiboly. The data for Nodal-overexpressing donor cells is also shown in (h). **(k)** Percentage of wt and Nodal-overexpressing donor cells, collected from 50% epiboly, shield or 75% epiboly stage embryos, which have arrived at the YSL by the end of MZ*oep* host embryo epiboly, as a function of the normalized intensity of nuclear pSmad2/3, averaged over the first 3 cell tiers for wt embryos and over the first 10 cell tiers for Nodal-overexpressing embryos (to account for the steep Nodal signalling gradient observed at the blastoderm margin of wt embryos and the homogeneous nuclear accumulation of pSmad2/3 observed in Nodal-overexpressing embryos, see Methods for details). The data plotted in this graph are also shown in Fig. 1i, 2e and in (d-f, h). Note that - consistent with the notion that Nodal signals regulate the window of competence for autonomous mesendoderm internalization - both Nodal signalling activity and cell internalization capacity decay during gastrulation in Nodal-overexpressing embryos. Moreover, Nodal-overexpressing donor cells, collected from the blastoderm margin of shield stage embryos, showed higher signalling activity than stage-matched wt embryos, but lower than wt marginal cells or Nodal-overexpressing cells in 50% epiboly stage embryos. Consequently, Nodal-overexpressing donor cells, collected from shield stage embryos, internalize more efficiently than stage-matched wt cells, yet less efficiently than wt marginal cells or Nodal-overexpressing cells from 50% epiboly stage embryos. This also supports the notion that Nodal signals regulate the autonomous internalization capacity of mesendoderm cells in parallel to cell fate^33^, since Nodal-overexpressing cells at all stages of gastrulation adopt an anterior axial mesoderm fate (a, b), but still show very distinct internalization capacities (h). **(l)** Length of protrusions formed by donor cells, collected from the blastoderm margin of wt and Nodal-overexpressing embryos control or coexpressing DN-Rac1, at 50% epiboly (wt: n=44 cells, N=7) or shield stage (wt: 36 cells, N=4; +Nodal: 47 cells, N=5; +Nodal+DN-Rac1: 48 cells, N=7), after transplantation into MZ*oep* hosts. The data shown for wt donor cells are also shown in Extended Data Fig. 2a. Each dot in the graph corresponds to the average length of cell protrusions in a single transplanted cluster at a given time point (each donor cell cluster was analysed every 5min for 60min, see Methods for details). Mann-Whitney test. \*\*\*\**P*<0.0001 (d-f), \**P*=0.0253 (j) (d-f, j). Kruskal-Wallis test. NS: not significant. \*\*\*\**P*<0.0001. \*\*\**P*=0.0003 (l). Dorsal view (top view: (a, b, c); cross-section: (g)); Side view (a,b). Scale bars: 100 µm (a, b), 20 µm (c, g).

**Extended Data Figure 4.**
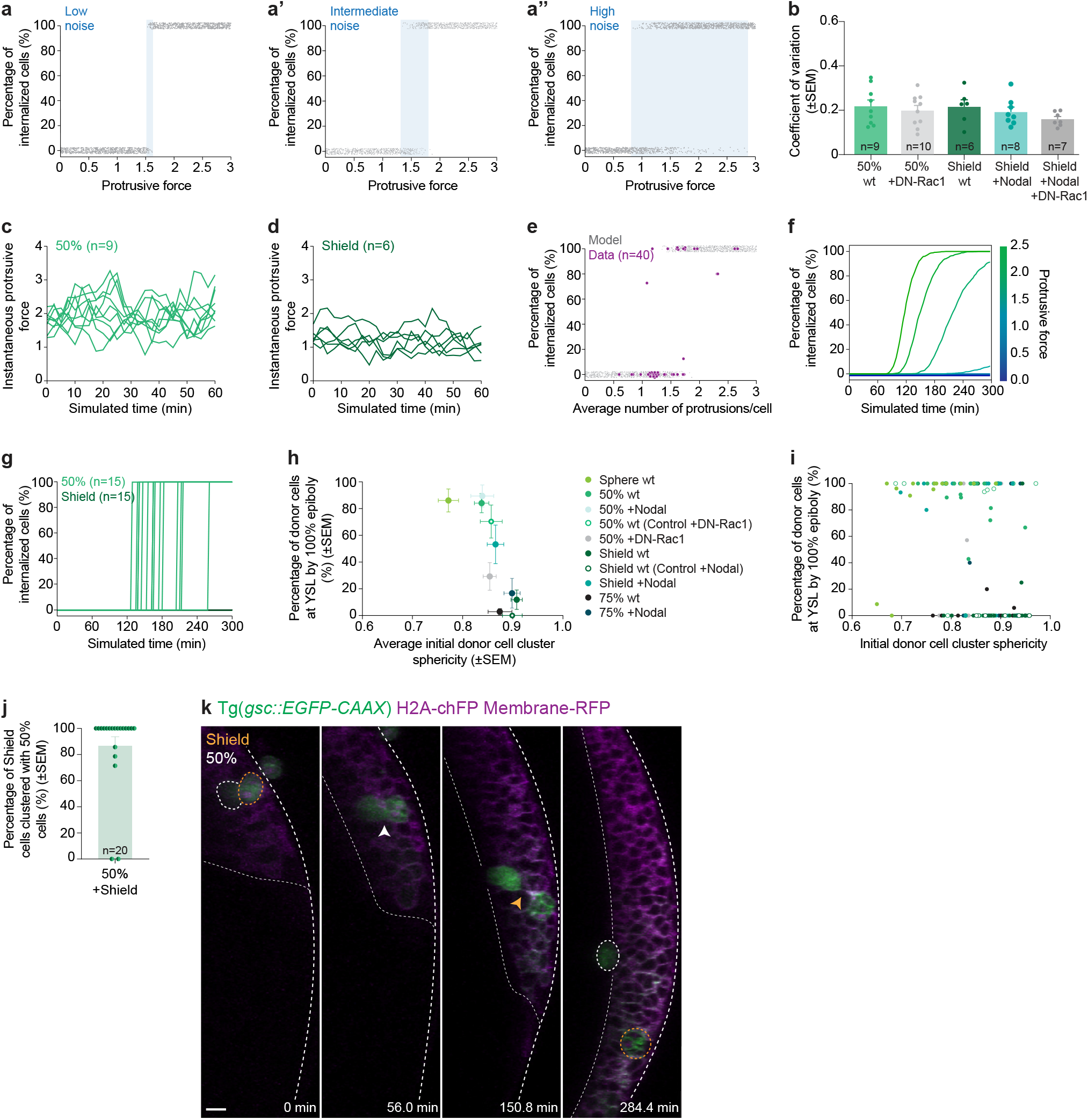
Motility-driven (un)jamming underlies the transition between leaders and follower cells, and contact between these distinct progenitors is required for follower cell internalization. **(a-a’’)** Prediction from the toy-model of motility-driven (un)jamming for the relationship between internalization capacity (accessed as the final position of cells by the end of a simulation) and average protrusive force *F* for different values of noise σ, corresponding to 5% (a), 20% (a’) or 100% (a’’) normalized standard deviation in the protrusive force (see Supplementary Note for details). Each grey dot represents the outcome of a simulation (n=2000 per panel; note that a small dispersion along the y axis was added to better convey the probability associated with a given internalization outcome as a function of *F*). Increasing noise σ in the protrusive force *F* results in a progressive increase of the bistable region (blue shaded interval), where for the same value of average protrusive force, cells can either internalize or stay out. **(b)** Coefficient of variation for the number of protrusions formed by control or DN-Rac1-expressing mesendoderm cells, collected from the blastoderm margin of wt and Nodal-overexpressing embryos at 50% epiboly (wt: 44 cells, N=7; +DN-Rac1: 54 cells, N=10) or shield stage (wt: 36 cells, N=4; +Nodal: 47 cells, N=5; +Nodal+DN-Rac1: 48 cells, N=7), after transplantation into MZ*oep* hosts. **(c, d)** Instantaneous protrusive forces for simulated donor cells clusters for average protrusive force *F*=1.86 (c, N=9) and *F*=1.15 (d, N=6), corresponding to the average number of protrusions formed by mesendoderm donor cells collected from 50% or shield stage embryos, respectively (see Fig. 2c and Extended Data Fig. 2b, c for comparison with the experimental data). **(e)** Percentage of internalized cells as a function of the average number of protrusions formed per cell across all experimental conditions and replicates (purple, each circle represents a single transplant - average data are shown in Fig. 3b) and predicted by the model (grey, same parameters as (a’), selected based on the experimentally observed variance in the cell protrusion number, as shown in (b), n=2000). The experimental data shown in the graph correspond to control or DN-Rac1-expressing donor mesendoderm cells, collected from the blastoderm margin of wt and Nodal-overexpressing embryos, at 50% epiboly (wt: 44 cells, N=7; +DN-Rac1: 54 cells, N=10) or shield stage (wt: 36 cells, N=4; +Nodal: 47 cells, N=5; +Nodal+DN-Rac1: 48 cells, N=7) and transplanted into MZ*oep* hosts (the average data is also shown in Fig. 3b). **(f)** Percentage of internalized cells as a function of time in the numerical simulations (same parameters as (a’)) for different values of protrusive force *F* (see colour-code, each curve corresponds to an increment of 0.35 in protrusive force: *F* = 0.35, 0.7, 1.05, 1.4, 1.75, 2.1, 2.45). **(g)** Percentage of internalized cells as a function of time in individual simulations (n=15 per condition), for protrusive force *F*=1.86 (light green) and *F*=1.15 (dark green), corresponding to the average number of protrusions formed by mesendoderm donor cells collected from 50% or shield stage embryos, respectively (see Fig. 2c). For *F*=1.15, we found no internalization events occur, while for *F*=1.86 internalization is robust, but highly asynchronous, mirroring the experimental findings shown in Extended Data Fig. 1c. **(h, i)** Percentage of control or DN-Rac1-expressing mesendoderm donor cells, collected from the blastoderm margin of wt and Nodal-overexpressing embryos at sphere (wt: n=15, N=6), 50% epiboly (wt: n=16, N=9; wt (+DN-Rac1 control): n=14, N=9; +DN-Rac1: n=19, N=9), shield (wt: n=19, N=12; wt (+Nodal control): n=7, N=4; +Nodal: n=12, N=4) or 75% epiboly stage (wt: n=9, N=8; +Nodal: n=12, N=6), which have arrived at the YSL of MZ*oep* hosts by the end of epiboly, as a function of their initial sphericity (see Methods for details). The average is shown in (h) and individual transplants in (i). **(j)** Percentage of co-transplanted mesendoderm donor cells, collected from the blastoderm margin of 50% epiboly and shield stage embryos and transplanted into MZ*oep* host embryos, which remain clustered until the hosts reached 100% epiboly stage (N=11). **(k)** High-resolution confocal images of exemplary co-transplanted mesendoderm donor cells, collected from the blastoderm margin of 50% epiboly and shield stage embryos and transplanted into MZ*oep* hosts. All donor cells express gsc::EGFP-CAAX (green) and can be distinguished by H2A-chFP expression (magenta, nuclei). Host embryos express low levels of gsc::EGFP-CAAX (green) and Membrane-RFP (magenta). Dashed white lines indicate EVL and YSL. White and yellow dashed lines outline donor cells collected from 50% epiboly and shield stage embryos, respectively. White arrowhead points at an initially cohesive co-transplanted heterotypic cluster. Yellow arrowhead indicates the separation of a heterotypic cluster, with early cells (50% epiboly) undergoing internalization and late cells (shield stage) remaining in more superficial regions of the blastoderm (n=2/20, N=10). Dorsal view (cross-section: (k)). Scale bar: 20 µm (k).

**Extended Data Figure 5.**
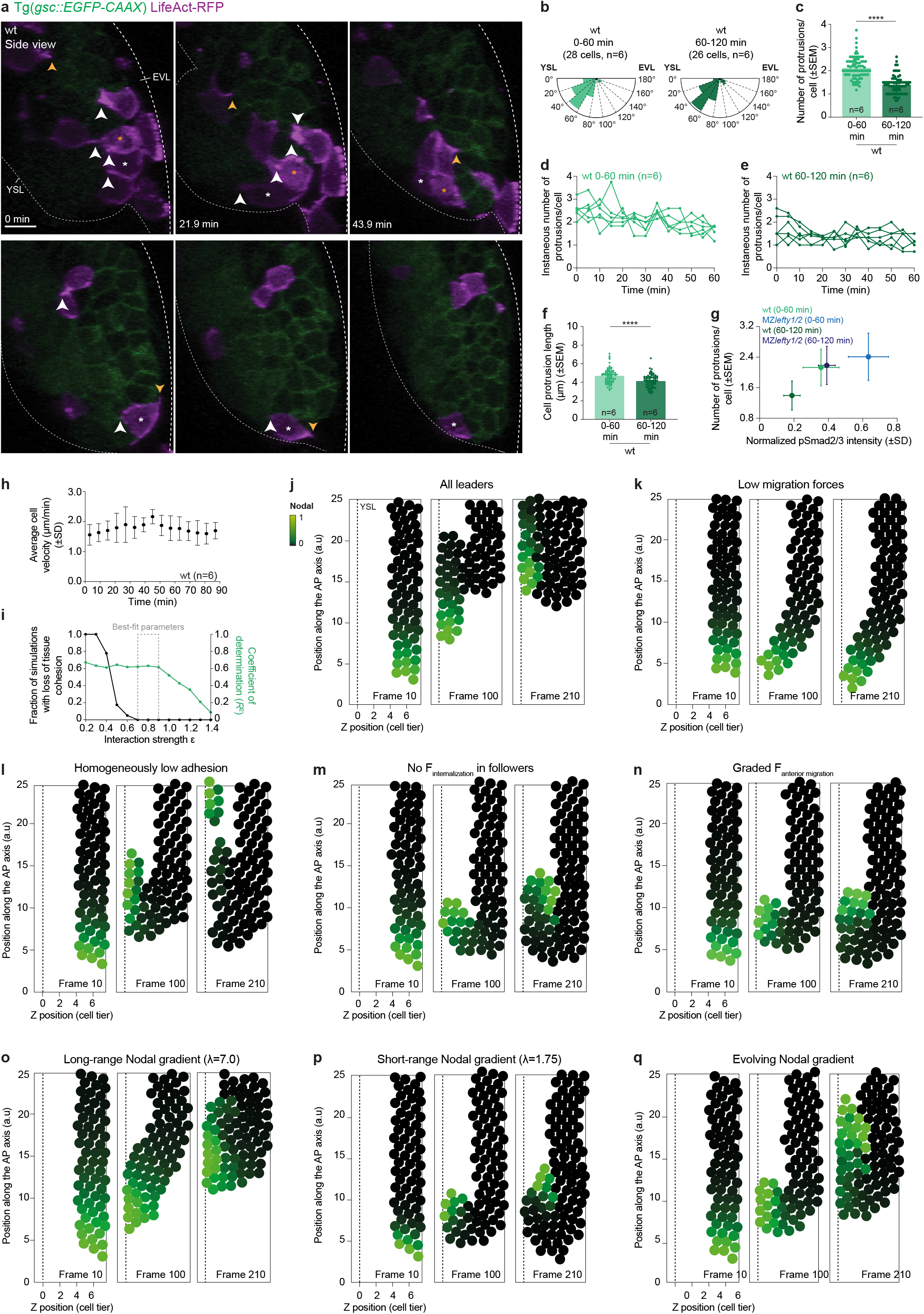
Decay of mesendoderm cell protrusiveness *in vivo* and sensitivity analysis of the numerical simulations. **(a)** High-resolution confocal images of an exemplary wt embryo during tissue internalization. Axial mesendoderm cells are marked by gsc::EGFP-CAAX expression (green), while actin-rich protrusions are labelled in a mosaic fashion by LifeAct-RFP (magenta). White and yellow arrowheads indicate cell protrusions oriented towards the YSL and EVL, respectively. Asterisks indicate internalizing cells, with the cell located closest to the blastoderm margin initiating internalization first (white asterisk top panel). Dashed white lines indicate the EVL and YSL. 0 min, internalization onset. **(b)** Rose plot of cell protrusion orientation in mesendoderm cells internalizing early (0-60 min: N=6) and late (60-120 min: N=6; see Methods for details). **(c)** Average number of protrusions formed by mesendoderm cells internalizing early (0-60 min: 28 cells, N=6) and late (60-120 min: 26 cells, N=6; see Methods for details). Each dot in the graph corresponds to the average number of protrusions formed per internalizing mesendoderm cell in a single embryo at a given time point (each donor cell cluster was analysed every 5min for 60min, see Methods for details). **(d, e)** Instantaneous number of protrusions formed per mesendoderm cell internalizing early (d, 0-60 min: 28 cells, N=6) and late (e, 0-120 min: 26 cells, N=6). Each curve corresponds to the average number of protrusions formed per internalizing mesendoderm cell in a single embryo (see Methods for details). **(f)** Length of protrusions formed by mesendoderm cells internalizing early (0-60 min: 28 cells, N=6) and late (60-120 min: 26 cells, N=6; see Methods for details). Each dot in the graph corresponds to the average length of cell protrusions formed by internalizing mesendoderm cells in a single embryo at a given time point (each donor cell cluster was analysed every 5min for 60min, see Methods for details). **(g)** Average number of protrusions formed by wt or MZ*lefty1/2* mesendoderm cells as a function of their normalized pSmad2/3 nuclear accumulation averaged over the first 3 cell tiers (see Methods for details). The data plotted in this graph are also shown in Fig. 2e, 4e, Extended Data Fig. 6b, j and in (c). **(h)** Mesendoderm cell velocity, averaged at each time points across all tracked wt embryos (N=6), during epiboly, internalization and anterior migration movements. 0 min, internalization onset. **(i)** Sensitivity analysis for different values of interaction strength ε and its impact on mesendoderm tissue cohesion and preservation of positional order in the numerical simulations. The dashed box indicates the range of best-fit parameters (ensuring both tissue cohesion and high R^2^) that are used in the simulations (see Supplementary Note for details). **(j-q)** Numerical simulations of mesendoderm cell internalization for different assumptions and parameter values. Particles are colour-coded for Nodal signalling activity. Mesendoderm cells are defined as any cell with a Nodal signalling >0.05 (see Supplementary Note for details). Dashed lines indicate the YSL. In (j), all mesendoderm cells, regardless of their Nodal signalling activity, were assumed to have equally high internalization forces. In this case, positional information during tissue internalization was lost, since mesendoderm cells located far from the margin reach the YSL boundary at the same time as cells positioned right at the margin. In (k), all parameters are unchanged from those shown in Fig. 4b (simulating the wt situation), except that lower migratory forces were assumed (both for internalization and animal migration forces). This resulted in a lack of large-scale internalization movements at the margin (turning motion), as migratory forces were insufficient to drive the cellular re-arrangements necessary for this morphogenetic movement. In (l), all parameters are unchanged from those shown in Fig. 4b, except that homogeneously lower adhesion between cells was assumed. This led to loss of tissue-level cohesion during internalization movements, suggesting that a minimum level of cell-cell adhesion is required for tissue integrity during internalization. In (m), all parameters are unchanged from those shown in Fig. 4b, except that zero internalization forces were assumed for all follower cells (defined as any cell with a Nodal signalling <0.5). In this case, internalization is similar to the situation shown in Fig. 4b, suggesting that the internalization forces of follower cells are largely dispensable to reproduce wt tissue internalization. In (n), all parameters are unchanged from those shown in Fig. 4b, except that, similarly to the internalization forces, the anterior migration-directed forces were assumed to be proportional to Nodal signalling activity (see Supplementary Note for details). While the overall tissue movements were, as expected, delayed compared to those shown in Fig. 4b, it still gave rise to ordered tissue internalization movement. In (o), all parameters are unchanged from those shown in Fig. 4b, except that an expanded Nodal gradient was assumed (see Supplementary Note for details). Similar to the phenotype of MZ*lefty1/2* embryos (Fig. 4g-k, Extended Data Fig. 6c and Supplementary Video 5), this led to the near-simultaneous internalization of many cells at the margin, thus resulting in loss of positional information. In (p), all parameters are unchanged from those shown in Fig. 4b, except that a shorter Nodal gradient was assumed (see Supplementary Note for details). This led to a smaller population of leader cells, which, nonetheless, was able to initiate robust and orderly tissue internalization. In (q), all parameters are unchanged from those shown in Fig. 4b, except that an evolving Nodal gradient was assumed (see Supplementary Note for details), whereby cells closest to the margin continue to increase Nodal signalling activity over time. This accelerates the process of internalization and anterior movements, while ordering is largely unchanged. Mann-Whitney test. \*\*\*\**P*<0.0001 (c, d). Dorsal view (cross-section: (a)). Scale bar: 20 µm (a).

**Extended Data Figure 6.**
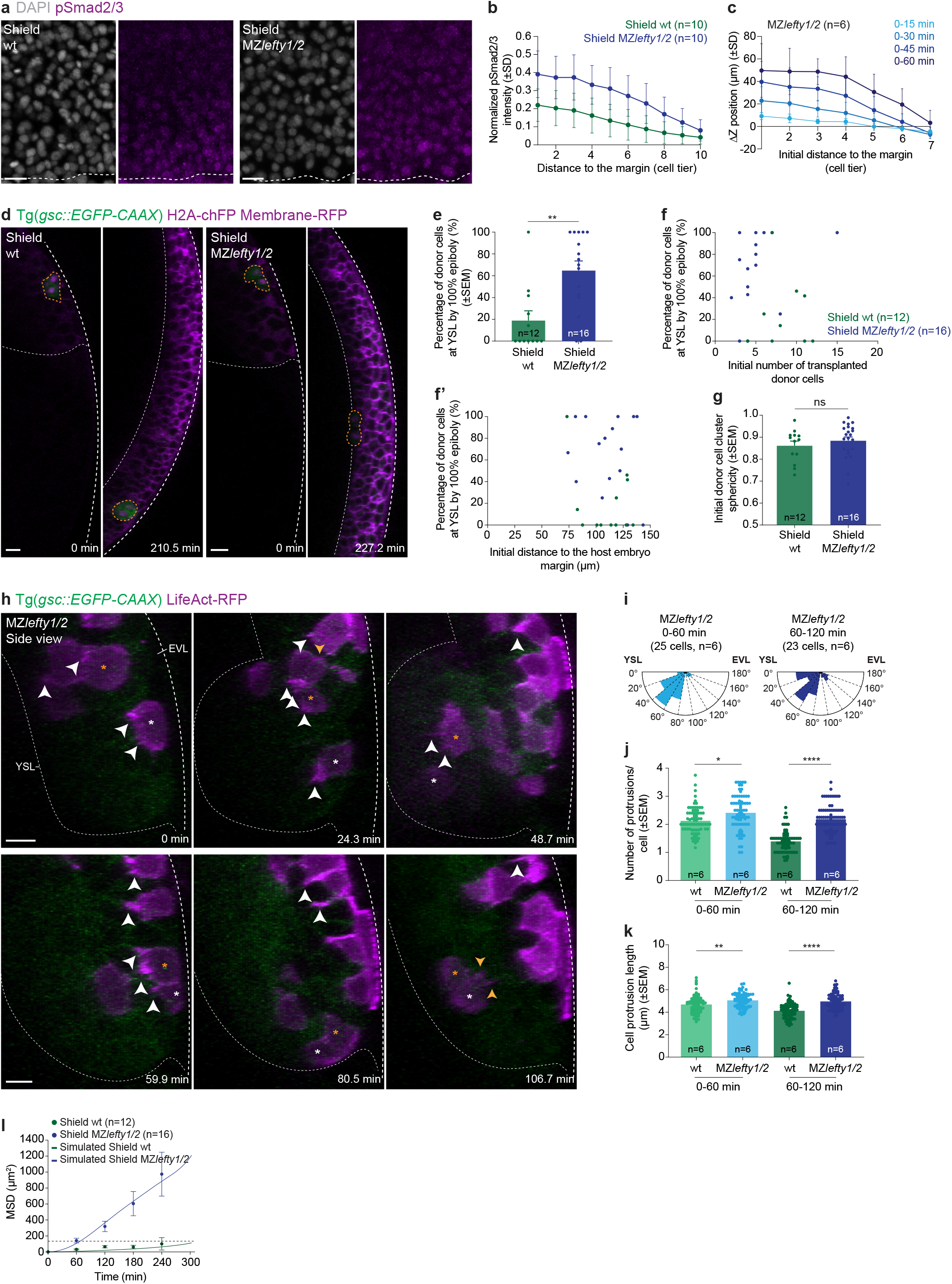
Expanding the Nodal signalling gradient affects the spatiotemporal pattern of leader-to-follower mesendoderm cells. **(a)** High-resolution confocal images of exemplary wt and MZ*lefty1/2* embryos stained for DAPI (grey, nuclei) and pSmad2/3 (magenta) at shield stage. Dashed lines indicate deep cell margin. **(b)** Normalized intensity of nuclear pSmad2/3 as a function of their distance to the blastoderm margin, expressed as cell tiers, in wt and MZ*lefty1/2* embryos at shield stage (N=3; see Methods for details). **(c)** Change in Z position, a proxy for cell internalization, of mesendoderm cells as a function of their initial distance to the margin, expressed in cell tiers, in MZ*lefty1/2* embryos (N=6). The colour-code corresponds to the indicated time bins. **(d)** High-resolution confocal images of exemplary wt and MZ*lefty1/2* donor cells, collected from the blastoderm margin of shield stage embryos and transplanted into MZ*oep* mutant host embryos. Donor cells are marked by gsc::EGFP-CAAX (green) and H2A-chFP expression (magenta, nuclei), while host embryos express low levels of gsc::EGFP-CAAX (green) and Membrane-RFP (magenta). For each transplant, the first acquired time point and the time point when the hosts reached 100% epiboly stage are shown. Dashed white lines indicate EVL and YSL; yellow dashed lines outline donor cell transplants. **(e)** Percentage of control (N=7) and MZ*lefty1/2* (N=6) mesendoderm donor cells, collected from the blastoderm margin of shield stage embryos, which have arrived at the YSL by the end of host embryo epiboly. **(f, f’)** Percentage of control (N=7) and MZ*lefty1/2* (N=6) mesendoderm donor cells, collected from the blastoderm margin of shield stage embryos, which have arrived at the YSL by the end of host embryo epiboly, as a function of the initial number of transplanted cells (f) or the initial distance of the transplanted cells to the host embryo margin (f’). **(g)** Initial sphericity of control (N=7) and MZ*lefty1/2* (N=6) mesendoderm donor cell clusters, collected from the blastoderm margin of shield stage embryos, after transplantation into MZ*oep* hosts. **(h)** High-resolution confocal images of an exemplary MZ*lefty1/2* embryo during mesendoderm internalization. Axial mesendoderm cells are marked by gsc::EGFP-CAAX expression (green), while actin-rich protrusions are labelled in a mosaic fashion by LifeAct-RFP (magenta). White and yellow arrowheads indicate cell protrusions oriented towards the YSL or EVL, respectively. Asterisks indicate internalizing cells. 0 min, internalization onset. Note that, in contrast to wt embryos, mesendoderm cells in MZ*lefty1/2* embryos positioned closer or further away from the blastoderm margin initiate internalization movements almost simultaneously, resulting in loss of positional information (compare white and orange asterisks in the top panel). **(i)** Rose plot of cell protrusion orientation in mesendoderm cells internalizing early (0-60 min: N=6) and late (60-120 min: N=6; see Methods for details) in MZ*lefty1/2* embryos. **(j)** Average number of protrusions formed by mesendoderm cells internalizing early (0-60 min) and late (60-120 min) in wt (28 and 26 cells, N=6) or MZ*lefty1/2* (25 and 23 cells, N=6) embryos (see Methods for details). Each dot in the graph corresponds to the average number of protrusions formed per internalizing mesendoderm cell in a single embryo at a given time point (each donor cell cluster was analysed every 5min for 60min, see Methods for details). **(k)** Length of protrusions formed by mesendoderm cells internalizing early (0-60) and later (60-120 min) in wt (28 and 26 cells, N=6) or MZ*lefty1/2* (25 and 23 cells, N=6) embryos (see Methods for details). Each dot in the graph corresponds to the average length of cell protrusions formed by internalizing mesendoderm cells in a single embryo at a given time point (each donor cell cluster was analysed every 5min for 60min, see Methods for details). The data shown for wt cells in (j, k) are also shown in Extended Data Fig. 5c and g. **(l)** Mean squared relative displacement of donor mesendoderm cells, collected from the blastoderm margin of wt (N=7) or MZ*lefty1/2* (N=6) embryos at shield stage, or predicted by the model based on their measured average protrusiveness (n=2000). The dashed line corresponds to the average cell size at 300 min (see Methods and Supplementary Note for details). The corresponding percentage of donor cells which have arrived at the YSL as a function of host developmental stage for these transplants is shown in (e) and the average number of cell protrusions formed per cell is shown in (j). Mann-Whitney test. \*\**P*=0.0027 (e). *t* test. NS: Not significant (g). Kruskal-Wallis test. *\*P*=0.0229 (j). *\*\*P*=0.0015 (k). *\*\*\*\*P*<0.0001 (j, k). Dorsal view (top view: (a); cross-section: (d, h)). Scale bars: 20 µm (a, d, h).

**Extended Data Figure 7.**
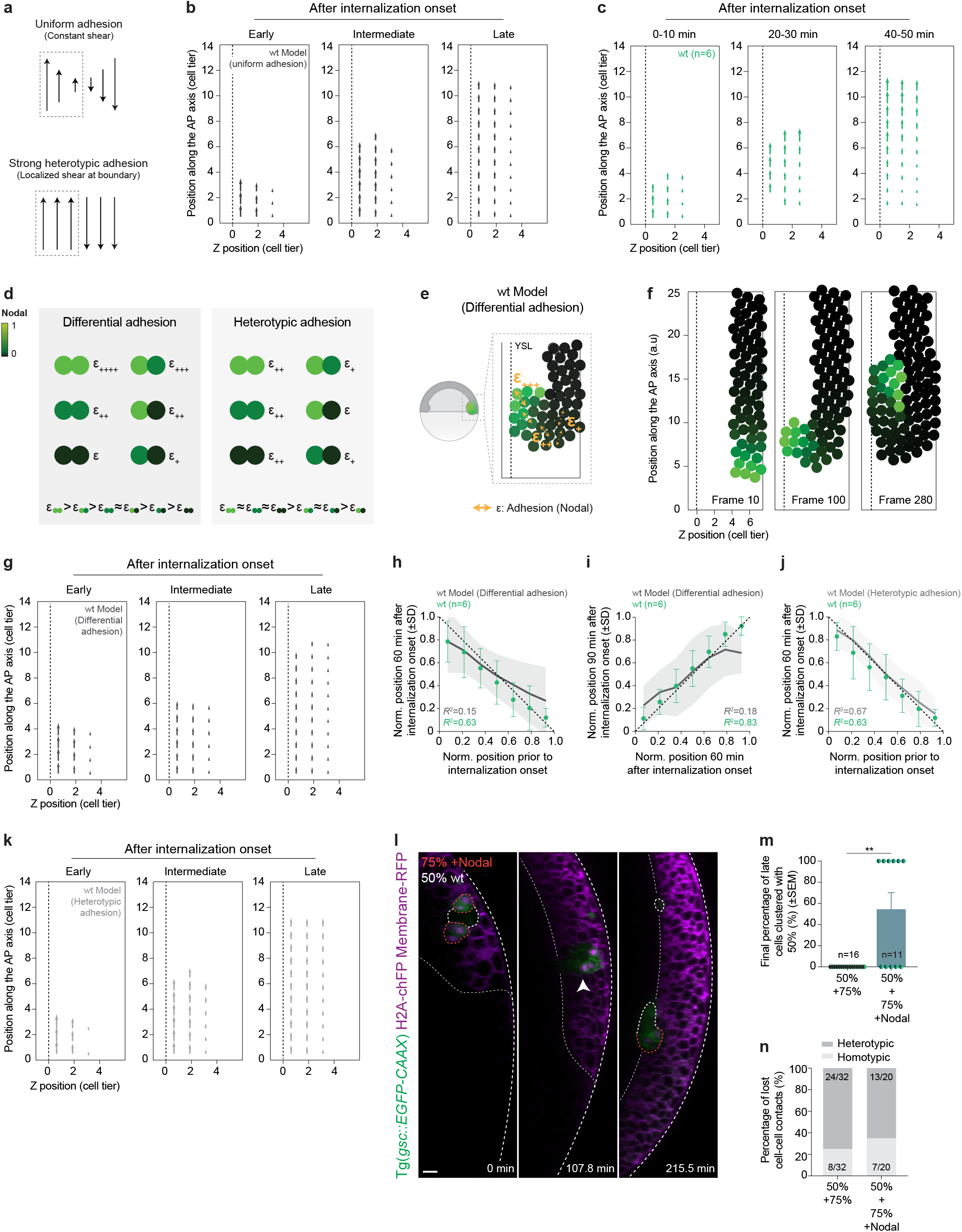
Modulating Nodal signalling in heterotypic clusters of transplanted mesendoderm cells is sufficient to rescue cluster splitting. **(a)** Schematic representation of the velocity gradients expected to arise at the boundary between internalized (highlighted with the grey dashed box) and non-internalized mesendoderm cells in the simulations for different model assumptions. In case of uniform cell-cell interactions, velocity gradients are expected to be constant, as observed in shearing of simple fluids (near-zero velocities arise at the boundary). In the case of strongly heterotypic interactions, velocity gradients are expected to be concentrated at the boundary (where interactions are weak), with near-constant velocities on either side. The dashed box highlights the velocity gradients in the internalized mesendoderm cells, the region we analysed in the next panels *in vivo* and i*n silico*. **(b)** Average velocity maps of internalized mesendoderm cells during anterior migration in wt simulations assuming uniform adhesion (see Supplementary Note for details). The maps were ‘stage-matched’ to the average position of internalized mesendoderm cells along the anterior-posterior axis *in vivo* (early, intermediate and late correspond to frames 80, 130, 250, respectively). Dashed lines indicate the YSL. **(c)** Average velocity maps of internalized mesendoderm cells during anterior migration in wt embryos (N=6, see Methods for details). Dashed line as in (b). **(d)** Schematic representation of the Nodal signalling-dependent differential and heterotypic/preferential adhesion models and their implications for the strength of contacts between cells with different Nodal signalling activity (see Supplementary Note for details). Particles are colour-coded for Nodal signalling activity. **(e)** Schematic representation of the 2D-particle based model used to simulate mesendoderm internalization assuming Nodal signalling-dependent differential adhesion (i.e. adhesion gradient, see Supplementary Note for details). All remaining parameters were left as in Fig. 4b. Colour-code as in (d) and dashed line as in (b). **(f)** Numerical simulations using the same parameters as in Fig. 4b, except that Nodal signalling-dependent differential, rather than uniform, adhesion was assumed (see Supplementary Note for details). Colour-code as in (d) and dashed lines as in (b). Similar to our simulations assuming uniform adhesion, in these simulations, leader cells are highly adhesive and, therefore, strongly interact with the overlying non-internalized cells, resulting in loss of positional information. **(g)** Average velocity maps of internalized mesendoderm cells during anterior migration in wt simulations assuming Nodal signalling-dependent differential adhesion (see Supplementary Note for details). The maps were ‘stage-matched’ to the average position of internalized mesendoderm cells along the anterior-posterior axis *in vivo* (early, intermediate and late correspond to frames 100, 150, 280, respectively). Dashed lines as in (b). **(h, i)** Correlation between cell position at early (h) and late (i) stages of tissue internalization in wt simulations assuming Nodal signalling-dependent differential adhesion (*R*^2^=0.15 (h), *R*^2^=0.18 (i), N=20 simulations) and wt embryos (*R*^2^=0.63 (h), *R*^2^=0.83 (i), N=6; see Methods and Supplementary Note for details). Dashed line indicates perfect conservation of the relative position of mesendoderm cells during internalization (*R*^2^=1). **(j)** Correlation between cell position at the onset of tissue internalization in wt simulations assuming Nodal signalling-dependent heterotypic/preferential adhesion (*R*^2^=0.67, N=20 simulations) and wt embryos (*R*^2^=0.63, N=6; see Methods and Supplementary Note for details). Dashed line as in (h, i). **(k)** Average velocity maps of internalized mesendoderm cells during anterior migration in wt simulations assuming Nodal signalling-dependent heterotypic interactions (see Supplementary Note for details). The maps were ‘stage-matched’ to the average position of internalized mesendoderm cells along the anterior-posterior axis *in vivo* (early, intermediate and late correspond to frames 70, 120, 210, respectively). Dashed lines as in (b). **(l)** High-resolution confocal images of an exemplary heterotypic cluster of co-transplanted mesendoderm cells, collected from the blastoderm margin of wt embryos at 50% epiboly stage and Nodal-overexpressing embryos at 75% epiboly stage, after transplantation into MZ*oep* hosts. All donor cells express *gsc::EGFP-CAAX* (green) and can be distinguished by H2A-chFP expression (magenta, nuclei). Host embryos express low levels of *gsc::EGFP-CAAX* (green) and Membrane-RFP (magenta). Dashed white lines indicate the EVL and YSL. White lines and orange dashed lines outline donor cells collected from wt embryos at 50% epiboly and Nodal-overexpressing embryos at 75% epiboly stage, respectively. White arrowhead points at a cohesive heterotypic cluster. **(m)** Percentage of co-transplanted control (N=9) or Nodal-overexpressing (N=7) donor cells collected from the blastoderm margin of 75% epiboly stage embryos, which remain clustered with early cells (50% epiboly stage) until the end of MZ*oep* hosts epiboly. **(n)** Percentage of homotypic and heterotypic cell-cell contacts lost upon the final splitting of mesendoderm donor cell clusters of different compositions (50%+75%: N=9; 50%+75% +Nodal: N=7) until the MZ*oep* hosts reached 100% epiboly stage. The data shown for heterotypic clusters composed of donor cells collected from wt embryos at 50% and 75% epiboly (50%+75%) stage in (m, n) are also shown in Fig. 5g, h. Mann-Whitney test. *\*\*P*=0.0016 (m). Dorsal view (cross-section: (l)). Scale bar: 20 µm (l).

## Supplementary videos

**Supplementary Video 1. Mesendoderm internalization in wt embryos.**

High-resolution confocal time-lapse of an exemplary wt embryo during mesendoderm internalization. Axial mesendoderm cells are marked by *gsc::EGFP-CAAX* expression (green), while H2A-chFP (magenta) marks the cell nuclei. Arrowheads indicate the relative positions of exemplary cells during internalization. Note that the cell highlighted with the grey arrow undergoes division and we subsequently indicate the only daughter cell visible in this cross-section during internalization. 0 min, internalization onset (see also Fig. 1b).

**Supplementary Video 2. Rapid decay in the internalization capacity of dorsal mesendoderm cells during gastrulation.**

High-resolution confocal time-lapse of exemplary wt mesendoderm donor cells, collected from the blastoderm margin of 50% epiboly, shield or 75% epiboly stage embryos, transplanted into MZ*oep* hosts. Donor cells are marked by *gsc::EGFP-CAAX* (green) and H2A-chFP expression (magenta, nuclei), while host embryos express low levels of *gsc::EGFP-CAAX* (green) and Membrane-RFP (magenta). The stage of donor and host embryos are indicated at the top (and shown schematically in Fig. 1g). Yellow dashed lines outline the donor cell transplants. 0 min corresponds to the first acquired time point after transplantation, and the last time point shown corresponds to the end of host embryos epiboly (see also Fig. 1h, i).

**Supplementary Video 3. Contact between leader and follower mesendoderm cells results in follower cell internalization.**

High-resolution confocal time-lapse of exemplary co-transplanted mesendoderm donor cells, collected from the blastoderm margin of 50% epiboly and shield stage embryos and transplanted into a MZ*oep* mutant host embryo. All donor cells express *gsc::EGFP-CAAX* (green) and can be distinguished by H2A-chFP expression (magenta, nuclei). Host embryos express low levels of *gsc::EGFP-CAAX* (green) and Membrane-RFP (magenta). White and yellow dashed lines outline donor cells collected from 50% epiboly and shield stage embryos, respectively. 0 min corresponds to the first acquired time point after transplantation and the last time point shown corresponds to the end of host embryo epiboly (see Fig. 3d-f).

**Supplementary Video 4. Numerical simulations of mesendoderm internalization.**

Numerical simulations of mesendoderm internalization, based on the experimentally-measured Nodal signalling gradient at the onset of gastrulation in wt or MZ*lefty1/2* embryos, assuming uniform adhesion or heterotypic/preferential adhesion (see also Fig. 4b, f, 5a, d and see Supplementary Note for details). Particles are colour-coded for Nodal signalling activity. Black lines indicate the YSL. The first frame shown in the numerical simulation is 3 in order to allow the initially noisy configuration to equilibrate and the frame rate is indicated at the bottom of the simulation box. The simulations were stopped once they reached the same average number of internalized mesendoderm cells observed *in vivo* (see Supplementary Note for details).

**Supplementary Video 5. Mesendoderm internalization in MZ*lefty1/2* embryos.**

High-resolution confocal time-lapse of an exemplary MZ*lefty1/2* embryo during mesendoderm internalization. Axial mesendoderm cells are marked by *gsc::EGFP-CAAX* expression (green), while H2A-chFP (magenta) marks the cell nuclei. Arrowheads indicate the relative positions of exemplary cells during internalization. Note that the cell highlighted with the dark green arrow undergoes division and we subsequently indicate the position of both daughter cells during internalization (when visible in this cross-section). 0 min, internalization onset (see also Fig. 4g).

**Supplementary Video 6. Loss of cluster cell cohesion at heterotypic cell-cell contacts.**

High-resolution confocal time-lapse of exemplary co-transplanted mesendoderm donor cells, collected from the blastoderm margin of 50% and 75% epiboly stage embryos and transplanted into MZ*oep* mutant host embryos. All donor cells express *gsc::EGFP-CAAX* (green) and can be distinguished by H2A-chFP expression (magenta, nuclei). Host embryos express low levels of *gsc::EGFP-CAAX* (green) and Membrane-RFP (magenta). White and orange dashed lines outline donor cells collected from 50% epiboly and 75% epiboly stage embryos, respectively. White arrowhead points at co-transplanted donor cells forming a cohesive heterotypic cluster. Yellow arrowhead indicates the separation of the heterotypic cluster, with early cells (50% epiboly) undergoing internalization and late cells (75% epiboly) remaining in more superficial regions of the blastoderm. 0 min corresponds to the first acquired time point after transplantation and the last time point shown corresponds to the end of host embryo epiboly (see also Fig. 5f, left panels).

Dorsal view (cross-section: 1-3, 5-6).

Scale bars: 20 µms (1-3, 5-6).

## Methods

### Fish lines and husbandry

Zebrafish (*Danio rerio*) maintenance and handling was performed as described^88^. The following strains were used in this study: wild type AB or ABxTL, Tg(*gsc::EGFP-CAAX*)^89^, MZ*oep*^47^, MZ*lefty1/2*^62^, MZ*oep*; Tg(*gsc::EGFP-CAAX*) and MZ*lefty1/2*; Tg(*gsc::EGFP-CAAX*). Mutant transgenic lines were generated by crossing the Tg(*gsc::EGFP-CAAX*) transgenic line with MZ*oep* or MZ*lefty1/2* mutants. Embryos were raised at 25-31°C in E3 or Danieau’s (58 mM NaCl, 0.7 mM KCl, 0.4 mM MgSO_4_, 0.6 mM Ca(NO_3_)_2_, 5 mM HEPES, pH 7.6) media and staged as previously described^90^.

### Molecular biology

To synthesize mRNA for the dominant-negative version of Rac1a (DN-Rac1), Gateway technology (Invitrogen)^91^ was used to create a pCS2 vector containing the mutated version of Rac1a. The coding sequence of *rac1a* (NCBI reference sequence: NM_199771.1) was amplified using specific primers with additional Gateway recombination arms (5’GGGGACAAGTTTGTACAAAAAAGCAGGCTTAATGCAGGCCATAAAGTGTG-3′ and 5′-GGGGACCACTTTGTACAAGAAAGCTGGGTATCACAGAAGGAGACATCTTCTC-3′) from cDNA library of sphere stage wild type Tübingen embryos. The PCR product was recombined with *pDONR221* (Lawson#208) and the resulting entry clone was used as a template to create the DN-Rac1a construct. Briefly, the Rac1a entry clone was amplified with specific primers (5’-GCTGTGGGAAAAAATTGCCTTCTGATCAG-3’ and 5’- CTGATCAGAAGGCAATTTTTTCCCACAGC-3’) and mutated at residue 17 to substitute a threonine by an asparagine (T17N) as previously described^92,93^, using site-directed mutagenesis. The introduction of the mutation was verified by sequencing, and the DN-Rac1a entry clone was recombined with *pCSDest2* (Lawson #444) and *p3E-polyA* (Chien#302) to create the vector used for mRNA synthesis (mMessage mMachine SP6 transcription kit, ThermoFisher).

### Embryo microinjections

mRNA synthesis was performed using the mMessage mMachine Kit (Ambion), and injections in 1- or 128-cell stage embryos were performed as described^88^. The following mRNAs were injected at 1-cell stage: 50-100 pg *H2A-chFP*^94^, 50-100 pg *Membrane-RFP*^95^, 75 pg *LifeAct-RFP*^96^ and 7.5 pg *DN-Rac1* (generated in this study). To constitutively activate Nodal signalling, donor embryos were injected with 100 pg *Ndr2-chFP*^97^ mRNA and 2 ng of *sox32* morpholino (5’- GCATCCGGTCGAGATACATGCTGTT-3’), as described previously^28^. To label F-actin in a mosaic fashion in wild type or MZ*lefty1/2* embryos, a single blastomere was injected at 128-cell stage with 7.5 pg *LifeAct-RFP*^96^.

### Cell transplantation assays

Donor and host embryos were manually dechorionated with watchmaker forceps in a glass dish with Danieau’s medium. Mesendoderm cells (clusters initially composed of 2-30 cells) were collected from the dorsal margin of donor embryos, marked by EGFP-CAAX expression in Tg(*gsc::EGFP-CAAX*) embryos, at different stages of gastrulation using a bevelled borosilicate needle (20 μm inner diameter with spike, Biomedical Instruments) connected to a syringe system mounted on a micromanipulator. Since donor embryos overexpressing Nodal ligands and/or DN-Rac1 showed aberrant morphogenesis, sibling embryos injected only with a fluorescence reporter - H2A-chFP or LifeAct-RFP - were used as staging controls. In Nodal-overexpressing embryos, donor cells were still collected from the margin. Donor cells were then transplanted into the dorsal margin, marked by low expression of EGFP-CAAX, in MZ*oep*; Tg(*gsc::EGFP-CAAX*) host embryos underneath the EVL (schematic representation of the transplantation strategy in Fig. 1g and Extended Data Fig. 1g). The developmental stage of donor embryos is indicated in the figure and, unless stated otherwise, mesendoderm cells were transplanted into sphere stage MZ*oep* host embryos.

In co-transplantation experiments of heterotypic clusters, mesendoderm cells were sequentially collected from the dorsal margin of donor embryos at different developmental stages into the same bevelled borosilicate needle and then transplanted into the dorsal margin of sphere stage MZ*oep*; Tg(*gsc::EGFP-CAAX*) host embryos underneath the EVL (schematic representation of the co-transplantation strategy in Fig. 3d and 5f). The exact order of collection of donor cells was purposely varied. In triple co-transplantations assays, mesendoderm cells were sequentially collected from 50%, shield and 75% epiboly stage embryos or 75%, shield and 50% epiboly stage embryos.

### Whole-mount immunofluorescence (WMIF)

Wild type, Nodal-overexpressing and MZ*lefty1/2* embryos were fixed at different stages of gastrulation (50% epiboly, shield and 75% epiboly stage) in 4% paraformaldehyde (in PBS) overnight at 4°C. After fixation, the samples were washed 3-4x in PBS, dechorionated using watchmaker forceps and then transferred into Methanol (100%) and stored at −20°C for a minimum of 2 h. Whole-mount immunofluorescence for α-pSmad2/3 was performed as described previously^55^. Briefly, embryos were washed 5x for 5 min in PBS+1% (w/v) Triton X-100 and then blocked at room temperature 2x for 1 h in blocking solution (PBS+1% Triton X-100+10% goat serum+1% DMSO). The α-pSmad2/3 antibody (Cell signalling, Clone D27F4, Cat#:8828, 1:1000) was prepared in fresh blocking solution and incubated overnight at 4°C. After incubation with the primary antibody, embryos were washed 5x for 10 min in PBS+1% Triton X-100 and 3x for 1 h in PBS+0.1% (w/v) Triton X-100. The secondary antibody (Alexa Fluor 546 Goat Anti-rabbit IgG, Thermo Fisher Scientific, Cat#:A-11010, 1:500) was prepared in blocking solution, and the samples were then incubated overnight at 4°C. To label cell nuclei, DAPI (1:1000) was added to the secondary antibody solution for 20 min at room temperature before the start of the last round of washes. Finally, samples were washed 2x for 5 min in PBS+1% Triton X-100 and 5x for 15-20 min in PBS+0.1% Triton X-100.

### Whole-mount *in situ* hybridization (WMISH)

Wild type and Nodal-overexpressing embryos were fixed at different stages of gastrulation (50% epiboly, shield and 75% epiboly stage) in 4% paraformaldehyde (in PBS) overnight at 4°C. After fixation, embryos were quickly washed 3-4x in PBS and dechorionated using watchmaker forceps. Embryos were then transferred into Methanol (100%) and stored at −20°C for a minimum of 2 h. *In situ* probes for *gsc* and *sox17* were synthetized from partial cDNA sequences using the MEGAscript T7 RNA polymerase kit (Thermo Fisher Scientific, Cat#:AM1334) with Roche digoxigenin (DIG)-modified nucleotides (Sigma-Aldrich, Cat#:11277073910). WMISH were performed as previously described^98^ for *gsc in situ* hybridization (Anti-DIG-AP Fab fragments, Roche, Cat#11093274910, 1:1000). For *sox17 in situ* hybridization, an extra proteinase K treatment (30 s; Invitrogen, Cat#:25530049) was performed and 5% dextran sulfate (Sigma-Aldrich, Cat#:31404) was added to the hybridization solution. The staining reaction was stopped simultaneously for wild type and Nodal-overexpressing embryos. After staining, the samples were cleared with 100% Ethanol and washed several times with PBS or PBS + 0.1% (w/v) Tween 20. The embryos were then mounted in a petri dish containing small 2% agarose wells (800 µm x 800 µm, Microtissues) in PBS and imaged using an Olympus SZX 12 stereo-microscope equipped with a QImaging Micropublisher 5.0 camera.

### Image acquisition

For whole-embryo single-plane imaging, dechorionated embryos were mounted in a glass-bottom dish containing 2% agarose wells and immobilized in 0.6-0.7% low melting point agarose with the dorsal side of embryo, marked by EGFP-CAAX expression in Tg(*gsc::EGFP-CAAX*) embryos, oriented laterally. Imaging was performed using a Nikon Eclipse inverted wide-field microscope equipped with a Nikon 10x/NA 0.3 PH1 air objective and a fluorescent light source (Lumencor).

For live imaging experiments, dechorionated embryos were mounted in petri dishes with 2% agarose wells and immobilized in 0.6-0.7% low melting point agarose with the dorsal side of embryo, marked by EGFP-CAAX expression in Tg(*gsc::EGFP-CAAX*) embryos, facing the objective. Imaging was performed using a LSM 880 or LSM 800 upright microscope equipped with a Zeiss Plan-Apochromat 20x/NA 1.0 water immersion objective. Immunostained embryos were mounted as described for live samples and imaged in PBS using a LSM 880 upright microscope equipped with a Zeiss Plan-Apochromat 20x/NA 1.0 water immersion objective. To be able to compare fluorescence intensity in immunostained samples across experimental replicates and conditions, all imaging conditions were kept similar.

For high-magnification analysis of mesendoderm cell internalization and protrusion formation, dechorionated embryos were mounted in a glass-bottom dish (35mm, MatTek Corporation, Cat#: P35G-1.5-14-C) and immobilized in 0.6-0.7% low melting point agarose with the dorsal side of embryo, marked by EGFP-CAAX expression in Tg(*gsc::EGFP-CAAX*) embryos, facing the objective. The embryos were then imaged using a Zeiss LSM880 inverted microscope equipped with a Zeiss Plan-Apochromat 40x/NA 1.2 water-immersion objective. In all cases, mounted embryos were imaged in Danieau’s media at 28.5°C.

### Data analysis

All image data analysis was performed using Fiji (NIH)^99^ and/or Bitplane Imaris.

### Analysis of endogenous mesendoderm cell internalization

To characterize the spatiotemporal dynamics of mesendoderm cell internalization in wild type and MZ*lefty1/2* embryos, the Imaris Spot plugin was used to track individual cells during internalization. The internalization onset (0 min) was defined as the time point immediately before a clear shift in the Z position of mesendoderm cells at the blastoderm margin became clearly detectable. At 0 min, 40 cells located at varying distances from the blastoderm margin in wild type and MZ*lefty1/2* embryos were selected and tracked for 60 min in 3D confocal stacks, including through cell division events. For wild type embryos, the tracking of mesendoderm cells in 3D confocal stacks was extended for up to 90 min after internalization onset in order to determine how efficiently positional information is preserved also at later stages of tissue internalization and anterior migration. The analysis time was ultimately restricted to 90 min after internalization onset to avoid confounding effects arising from dorsal mesoderm convergence movements. Reference spots were manually added at the blastoderm margin, so that all embryos could be rotated into a similar orientation and 6 embryos/genotype were analysed. The position of all tracked cells as a function of time was then exported into an excel sheet and processed using a custom-made code.

To calculate the correlation analysis between initial (0 min) and final position (60 min), the relative position *x*_*i*_ (resp. *x*_*f*_) of cells along the *x*→ axis pre- and post-internalization was normalized so that *x* = 0 refers to the cell closest to the margin and x = 1 to the cell furthest away from it. Note that when a cell divided, *x*_*f*_ was defined as the average coordinate of the daughter cells at 60 min. For this analysis, we considered only cells that had started to internalize by 60 min, defined as any cell that moved, at least, by 15 µm towards the YSL. Thus, for ordered tissue internalization (first internalized cells at 0 min should be have migrated furthest away from the margin by 60 min), *x*_*i*_ and *x*_*f*_ would be expected to be perfectly anti-correlated (1-x slope, shown in dashed line in Fig. 1f for example). The correlation analysis between 60 and 90 min was performed in the same way, although in this case, as this analysis was performed on cells that had already internalized by 60 min, perfect order entailed a perfect correlation between *x*_*i*_ and *x*_*f*_ (x slope, show in dashed line in Fig. 5b for example).

To determine the time of internalization onset, the temporal evolution of the coordinate *z*_*j*_(*t*) for each cell *j* (i.e. how much a cell moves towards the YSL) was fitted. The profiles *t* were found to be best-fitted by the simple logistic function:

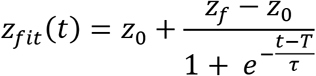

where *z*_0_ and *z*_*f*_ are fitting parameters denoting the initial (*z*_0_) and final (*z*_*f*_) z position, *T* is the time at which the internalization dynamics is maximum and *τ* is the characteristic duration of cell internalization. *t*_*in*_ = *T* − 2*τ* was then defined as the internalization onset for a given cell *j* and correlated to its initial distance *δx*_*j*_ to the margin in the *x* direction at *t=0*. This analysis showed that in wild type embryos, the onset of cell internalization *t*_*in*_ linearly correlated with the initial distance of mesendoderm cells to the margin (Fig. 1e), while in MZ*lefty1/2* embryos, this correlation was much weaker, since the first 2-4 cell rows were found to internalize almost simultaneously (Fig. 4j).

Finally, the average distance that a cell had moved in the z direction (*z*_*j*_(*t*) − *z*_*j*_(*t* = 0) for t=15, 30, 45 and 60 min), independently of any fitting procedure, was also quantified and correlated to its initial distance *δx*_*j*_ to the margin in the *x* direction at *t=0*. This initial distance to the margin *δx*_*j*_ was then binned according to the average cell size and averaged across 6 independent wild type and MZ*lefty1/2* embryos to show the mean and error bars (Extended Data Fig. 1b and 6c). To calculate the average cell size, the cell height of 20 cells/embryo was measured during early stages of tissue internalization.

### Analysis of mesendoderm donor cell internalization

To determine the internalization capacity of transplanted mesendoderm cells, the Spot plugin of Imaris was used to obtain the initial number and coordinates of donor cell nuclei, as well as reference landmarks at the blastoderm margin. Using a custom-made Matlab code^55^, all nuclei and reference landmarks were projected along the Z axis. The geometrical distance of the projected nuclei to the closest reference spot was then automatically calculated and exported along with their unique ID into an output excel sheet. Only donor cells placed within 150 µm to the blastoderm margin of MZ*oep* mutant host embryos were considered for subsequent analysis. A few additional criteria were considered for this analysis, namely (i) donor cell integrity must be preserved, at least, until host embryos reached 100% epiboly; (ii) donor cells must express comparable levels of EGFP-CAAX within a given transplanted cluster, to exclude cases where mesendoderm cells were co-transplanted with ectoderm or EVL cells; and, (iii) donor cells co-transplanted with yolk granules were removed from subsequent analysis. Similar exclusion criteria were employed for all the transplantation assays included in this study.

Using the same spot function aided by cross-section projections, the total number of transplanted cells and the number of donor cells which have arrived at the YSL were determined as a function of the developmental stage of the host embryo in order to normalize the data across embryos and experimental replicates. To determine the developmental stage of the host embryos, the Imaris Measurement tool was used to quantify the total embryo length, as well as the length covered by the deep cells at any given time point. This measurement was performed in a single-plane image at the bottom of the imaged volume, to be as close as possible to the centre of the embryo and then calculated as the percentage of deep cell epiboly. The percentage of mesendoderm donor cells, which have arrived at the YSL by the end of host embryo epiboly, were also plotted as a function of the (i) initial number of transplanted cells and (ii) initial distance of the transplanted cells to the host embryo margin.

### Analysis of mesendoderm cell protrusiveness

To analyse mesendoderm cell protrusion number, length and orientation, the multi-point tool of Fiji was used to obtain the coordinates at the base of each cell protrusion and at its tip. Using these vectors (base to tip), the orientation and length of cell protrusions were then calculated by translating the vectors in spherical coordinates *r*, *θ*, *φ*, using the classical physics convention.

Cell protrusion length:

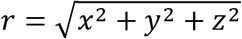

Angle *θ:*

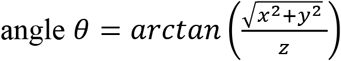

The angle *θ* quantifies the orientation of protrusions along the EVL-YSL axis. Protrusions are oriented exactly towards the EVL when *θ* = *π*, while protrusions are oriented exactly towards the YSL when *θ* = 0. F-actin-positive extensions exhibiting dynamic behaviour (appearance and eventual retraction), as well as blebs, defined as dynamic membrane extensions devoid of an F-actin rich cortex, were considered as cell protrusions and included in the analysis. We restrained from further categorizing these cellular protrusions, due to the complex 3D environment *in vivo*. Cell protrusions were analysed in 3D confocal stacks every 5 min for 1 h, either upon cell transplantation or within the intact embryo. The total number of measured cell protrusions was divided by the number of analysed cells to obtain the number of protrusions per cell. Mesendoderm cell protrusion number, length and orientation were averaged across time points and experimental replicates. The instantaneous fluctuation in the number of protrusions per cell is also shown in Extended Data Fig. 2c, d and 5d, e.

To relate cell protrusiveness to the internalization competence of mesendoderm cells, cell protrusions were analysed for the first h after the start of acquisition and the number of donor cells which have arrived at the YSL was determined 4 h later. Since these embryos were imaged at higher-magnification, the exact developmental stage of host embryos 4 h after the start of acquisition could not be determined, but likely corresponds to 90-100% epiboly stage. Notably, most of these transplantation assays were also repeated and imaged at lower resolution, confirming our observations in the high-magnification videos (the data is provided in Extended Data Fig. 2e-g’ and 3g-k). Since Nodal and/or DN-Rac1-overexpressing donor embryos displayed aberrant morphogenesis, sibling control embryos, injected with LifeAct-RFP alone, were used as staging controls.

To analyse mesendoderm protrusiveness in wild type and MZ*lefty1/2* embryos, two time windows - early (0-60 min) and late (60-120 min) - were defined to compare mesendoderm cell protrusion number, length and orientation. Internalization onset (0 min) was defined as the time point immediately before a clear shift in the Z position of mesendoderm cells at the blastoderm margin was detected. This temporal subdivision was chosen to compare the protrusiveness of mesendoderm cells internalizing early (‘leaders’) versus later (‘followers’) and corresponds approximately to the time difference between 50% epiboly and mid-to-late shield stage used for the cell transplantation assays (Extended Data Fig. 1a). For each temporal window, 3-5 cells were initially selected per embryo and both daughter cells were analysed in case of cell division.

### Analysis of the mean squared displacement (MSD) of mesendoderm donor cells

To compute the MSD of mesendoderm donor cell clusters upon transplantation into MZ*oep* hosts, we first measured the distance of each donor cell relative to the EVL, using a cross-section view and the Measurement tool in Imaris, every h until 100% host epiboly. We then averaged the distance of the cluster’s centre of mass to the EVL (proportional to *x*(*t*) in the toy-model of motility-driven (un)jamming, see Supplementary Note for details) and calculated the MSD (Extended Data Fig. 2d, 6l and Fig. 3c) as:

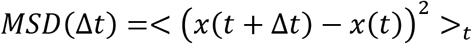

where <>_*t,n*_ denotes an average over all time points *t*. We only included time intervals (Δ*t*) for which we had, at least, n=5 transplants (up to 300min). To assess caged motion (i.e. MSD smaller than cell size on long time scales), we also determined the average cell size at 300 min (dashed line in Extended Data Fig. 2d, 6l and Fig. 3c), by measuring the cell height of 10 cells in 10 randomly selected host embryos.

### Analysis of mesendoderm donor cell sphericity

The degree of sphericity of a given cell cluster is controlled by the balance of adhesion forces at cell-cell contacts and cortical tension. This dimensionless parameter approaches 1 when the contact angle at cell-cell interface is close to 180° and, it progressively decreases with the reduction in this contact angle^58^. To analyse the initial sphericity of transplanted mesendoderm cell clusters, the number of transplanted cells and their initial distance to the host embryo margin was determined as described above. The Surface plugin of Imaris was then used to automatically segment the surface of the transplanted mesendoderm cell clusters, based on their EGFP-CAAX expression. In mesendoderm donor cells collected from sphere stage Tg(*gsc::EGFP-CAAX*) embryos, where *gsc::EGFP-CAAX* expression was still low, the outline of the cell clusters along their Z axis was, in some cases, manually determined to obtain an appropriate surface segmentation. In cases where there was more than one transplanted donor cell cluster initially, the same analysis was performed for each individual cluster and subsequently averaged across all transplants for the average analysis shown in Extended Data Fig. 4h and average per transplant for the analysis shown in Extended Data Fig. 4i.

### Analysis of pSmad2/3 nuclear localization

Analysis of the nuclear accumulation of pSmad2/3 was performed as described^55^. In short, the Imaris Spot detection plugin was used to obtain the 3D nuclear coordinates of all cells, based on their DAPI signal. The nuclei of EVL cells, cells undergoing division and cells displaying low intensity DAPI-positive nuclei were then manually excluded from the subsequent analysis. The remaining nuclei were filtered for (i) Z position, in order to restrict the analysis to the top 100 µm and avoid depth-related intensity changes, and (ii) maximum fluorescence intensity for both pSmad2/3 and DAPI below 65536 grey values to avoid oversaturated pixels. For shield and 75% epiboly stage embryos, we restricted our analysis to non-internalized mesendoderm progenitors by manually removing the nuclei of already internalized cells, aided by cross-section projections.To determine the distance of a given nuclei to the blastoderm margin, reference spots at the edge of the deep cell margin were manually added. Using a custom-made Matlab code^55^, the nuclei were projected along the Z axis. The geometrical distance of projected nuclei to the closest reference spot was then automatically calculated and exported along with their unique ID and mean fluorescence intensity for both pSmad2/3 and DAPI channels. For background subtraction, 15 spots were manually added within the cytoplasm of cells located at the bottom of the imaged volume. To try to correct depth-related artefacts, the normalized intensity of pSmad2/3 was calculated as the ratio between the mean fluorescence intensities for nuclear pSmad2/3 and DAPI, after background subtraction for both channels.

To plot the distance of a given nuclei to the blastoderm margin as cell tiers, we first measured the average size of 10-15 cells per wild-type embryo at 50% epiboly, shield and 75% epiboly stage. For each developmental stage, we then subtracted the minimum distance obtained between any nuclei and the reference spots and binned the data according with the measured cell size. The normalized intensity of nuclear pSmad2/3 as a function of their distance to the margin was then averaged across experimental replicates.

### Analysis of heterotypic mesendoderm donor cell clusters

To analyse the cohesion of heterotypic mesendoderm donor cell clusters after transplantation into MZ*oep* hosts, a proxy for functional differences in cell-cell adhesion strength, the initial composition of the heterotypic cluster and the distance of all donor cells to the blastoderm margin were determined as described above. By the end of epiboly in host embryos, the Spot plugin of Imaris was used to quantify the total number of ‘late’ mesendoderm donor cells (collected from shield, wild type or Nodal-overexpressing 75% epiboly, or shield+75% epiboly stage donor embryos) that remained clustered with ‘early’ cells (collected from 50% epiboly stage embryos). For heterotypic clusters composed of donor mesendoderm cells collected from the blastoderm margin of 50% and shield stage embryos, which tend to remain cohesive by the end of host embryo epiboly (Extended Data Fig. 4j), the proportion of ‘late’ shield cells which had arrived at the YSL was also quantified as described above (Fig. 3e, f). In cases where heterotypic clusters (partially or completely) split, the type of cell-cell contact (homotypic versus heterotypic) lost upon cluster splitting were recorded. A contact loss within the transplanted cluster was defined as a splitting event in case the split lasted more than 2 consecutive frames (approximately 10-15 min).

### Velocity maps for internalized mesendoderm cells

To analyse the dynamics of internalized mesendoderm cells during anterior migration, the Imaris Spot plugin was used to track individual cells, located at different Z positions relative to the YSL, in 3D confocal stacks. Here, 0 min corresponds to the time point during mesendoderm internalization where the first internalized mesendoderm cells reached the YSL. Cells were tracked for 10 min time intervals (0-10, 20-30 and 40-50 min) to obtain their instantaneous speed as a function of their distance to the YSL and blastoderm margin. Dividing cells were purposely excluded from tracking, since divisions are likely to delay the instantaneous speed of migrating cells. The velocity maps were restricted to the first hour of anterior migration to avoid confounding effects arising from dorsal mesoderm convergence movements. This procedure was then repeated for 6 independent embryos and the positions of all tracked cells as a function of time were recorded and processed using a custom-made code.

The instantaneous cell velocity was spatially binned both along the X (distance to the blastoderm margin) and Z axis (distance to the YSL), with bin size corresponding to the average cell width/height. To calculate the average cell size, the cell height of 20 cells/embryo in each time bin was measured. To normalize the XZ coordinates of each embryo, and be able to average across embryos, reference landmarks were manually added along the blastoderm margin and the newly formed tissue boundary at the start of each time bin using the Imaris Spot plugin. These reference coordinates were then used to calculate the geometrical distance of the cell nuclei to the blastoderm margin and the YSL and average the instantaneous speed along both axes in spatial bins. The embryos were also computationally rotated along the XZ plane to ensure that the average Z velocity was zero. This corrected for the overall curvature of the embryo and the newly formed tissue boundary, and also normalized the distance of a given cell nuclei to the margin across experimental replicates. A minimum of 3 tracks per bin/per sample were used for a given spatial and temporal bin to be plotted in the final velocity maps shown in Extended Data Fig. 7c.

### Statistics

The statistical analyses and plots were performed using Microsoft Excel, Gnuplot and/or GraphPad Prism. The number of transplants or embryos (n) and experimental replicates (N) analysed are indicated in the figure legends. At least three independent experiments were performed in all cases, except for the *sox17 in situ* hybridization shown in Extended Data Fig. 3b, where only two independent experiments were conducted. Error bars are indicated in the figures. No statistical test was performed to determine the sample sizes. The statistical tests used to access significance are indicated in the figure legends along with the corresponding *p* values. To choose an appropriate statistical test, the distribution of each group was first analysed using the D’Agostino-Pearson normality test. To compare two groups, a two-tailed Student *t* test or a Mann-Whitney test was used, depending on whether either dataset shows normal distribution. In cases where a *t*-test was used, the variances were subsequently tested using the F-test. To compare more than two groups, either ANOVA or Kruskal-Wallis test were performed, depending on whether all groups show normal distribution. To increase statistical power, a correction for multiple comparisons was used in these cases.

