## Supplementary Theory Note for "Morphogen gradient orchestrates pattern-preserving tissue morphogenesis via motility-driven (un)jamming"

**Supplementary Note**

Toy-model of motility-driven (un)jamming with directed polarity

Previous modelling approaches showed that motility-driven (un)jamming can arise from random motility forces providing active fluctuations able to overcome the local energy barriers linked to cell-cell rearrangements in solid tissues[1]. Such energy barriers can be modelled in a simplified manner as donor cell clusters at a position $x$ experiencing a bistable energy potential:

$$V_B(x) = \epsilon_b(-\frac{x^2}{2} + x^4)$$

where $x = 0.5$ and $x = -0.5$ correspond to the two stable states and $\epsilon_b$ is an energy scale setting the height of the potential (Fig. 3a). We thus model the resistance of the host tissue via this potential, denoting $x = -0.5$ as the initial position of cells (i.e. close to the EVL) and $x = 0.5$ the position corresponding to the inside of the host (i.e. close to the YSL). The other two forces acting on donor cell clusters are respectively active migratory forces $F(t)$ and a friction force, proportional to cluster velocity with respect to the host tissue $-\zeta\dot{x}(t)$, where $\zeta$ is a friction coefficient and $\dot{x}(t)$ the cluster velocity as a function time. The force balance therefore reads:

$$-\zeta\dot{x}(t) + F(t) - \partial_x V_b(x) = 0$$

Finally, we must complement this with a description of the motility force $F(t)$ itself. In contrast to the previous model of motility-driven (un)jamming, motility forces are not isotropic in this system, as we found that mesendoderm cells are polarized towards the YSL in all conditions considered (Fig. 2b, g, Extended Data Fig. 5b and 6i). The simplest description is then to take $F(t) = F_0$ as a constant for a given donor cell transplant, inferred from the experimentally measured number of protrusions per cell for a given experimental condition (Fig. 2c, h, Extended Data Fig. 5c and 6j). In this case, the dynamics is purely deterministic, and there is a critical force $F_c = \epsilon_b/3\sqrt{3}$ above which all donor cell clusters are predicted to internalize (with typical velocity $F_0/\zeta$), and, below which no clusters would be expected to internalize. In the presence of stochastic noise on the motility force $F(t)$, however, one expects that noise might also contribute for donor cell cluster internalization. To model this, we use the classical description of cell migratory forces $F(t)$ as an Ornstein-Uhlenbeck process:

$$dF(t) = \frac{F_0 - F(t)}{\tau_p} dt + F_0 \sigma dW_t$$

where $dW_t$ denotes the Wiener process, so that $F(t)$ fluctuate around a constant value $F_0$ (dependent on Nodal signalling activity in mesendoderm cells, Extended Data Fig. 5g) with persistence time $\tau_p$ and noise strength $\sigma$. In this model, the predicted variance in migratory forces can be calculated as $s = \frac{\sigma^2 F_0^2 \tau_p}{2}$, or alternatively the relative standard deviation $RSV = \sigma\sqrt{\frac{\tau_p}{2}}$. Since we found that the standard deviation of donor cell protrusiveness in time is consistently around 20% of the average ($RSV \approx 20\%$, Extended Data Fig. 4b-d), we included an $F_0$ coefficient in the noise term. Other sources of noise could in principle be considered (e.g. heterogeneity in the resistance of the host tissue), however we concentrated here on temporal stochasticity in donor cell protrusiveness as a minimal modelling assumption sufficient to explain our dataset (see below). Although, in the model, varying $F_0$ and $\tau_p$ independently, while constraining the resulting variance $s$ to the experimentally-measured value (Extended Data Fig. 4b), did not yield substantially different results, we could also constrain the value of $\tau_p$ by measuring the autocorrelation function of donor cell protrusiveness in time experimentally. Indeed, this is expected to decay with characteristic time scale $\tau_p$, leading us to estimate $\tau_p \approx 10\ min$, a value in line with measurements from other systems[2]. Once this is fixed, there is only a single other parameter, set by friction $\zeta$, controlling the timescale of internalization compared to the total simulation time $T_{tot}$ which we take as $\frac{\zeta}{T_{tot}} = 0.5$ so that cells above the unjamming threshold have internalized at the end of the numerical simulation (Extended Data Fig. 4f). Overall, the full predicted dynamics then only depends on a limited number of dimensionless parameters: $F_0/\epsilon_b$, the rescaled average protrusive/motility forces and $\sigma$, the noise associated with protrusion formation over time.

We thus performed a parameter screen for different values of these two parameters (n=2000 simulations per parameter value). As expected, for low values of noise $\sigma$, we recovered the previous deterministic limit, with a single well-defined protrusion force $F_c$ controlling a sharp transition from 0% to 100% internalization capacity (Extended Data Fig. 4a). Furthermore, increasing noise $\sigma$ gradually smoothed this transition by creating an intermediary region of the phase diagram (around the "deterministic critical force" at zero noise $F_c$), where bistable outcomes are observed (meaning that for the same average $F_0$, some simulated clusters will be internalized, while others remain out; Extended Data Fig. 4a-a'').

This is expected, as noise can allow donor cell clusters to internalize stochastically despite their average protrusiveness being slightly below the deterministic threshold. Importantly, when inputting the value of noise $\sigma$ corresponding to the experimentally measured value in protrusiveness variance (20% of the average, Extended Data Fig. 4b-d), we obtained a region of bistable outcomes of similar magnitude to what we found experimentally observed across all conditions (Extended Data Fig. 4e). When comparing the predicted relationship between average internalization capacity and protrusion force $F_0$ with the data averaged for each of the experimental conditions examined, we obtained an excellent qualitative and quantitative agreement (Fig. 3b). Thus, this minimal modelling approach argues that the degree of directed donor cell protrusiveness can be the control parameter for a motility-driven (un)jamming transition *in vivo*.

Finally, we further tested this toy-model using more detailed dynamical measurements of donor cell cluster internalization, by concentrating on the initial transplantation assays (donor cells collected from the dorsal margin of 50% epiboly and shield stage embryos, Fig. 1g-i, Extended Data Fig. 1c-e and Supplementary Video 2). Theoretically, a key signature of donor cell jamming is not simply a slower rate of displacement within the host tissue, compared to unjammed cells (i.e. internalization-competent cells), but that these cells should be "caged", i.e. not be able to move more than one cell diameter even over long time scales. We thus computed the mean squared displacement (MSD) of donor cell cluster position as a function of time (see Methods for details). We also computed the average cell size $l$ by 300 min, the last analysed time point, so that a MSD plateauing below $l^2$ at long time scales demonstrates "caged" motion. Indeed, we found that the MSD of donor cell clusters collected from the dorsal margin of shield stage embryos (i.e. internalization-incompetent cells, Fig. 1h, i and Supplementary Video 2) displayed a characteristic caged plateau even at long time scales (300 min), while the MSD of donor cells clusters collected from 50% epiboly stage embryos (i.e. internalization-competent cells, Fig. 1h, i and Supplementary Video 2) grew continuously in time (Extended Data Fig. 2d). Importantly, we found good qualitative and quantitative agreement when comparing these experimental measurements to our previously calibrated model (Fig. 3c). We also extended this analysis to mesendoderm donor cell clusters collected from MZ*lefty1/2* embryos at shield stage and, again, found a good agreement between the experimentally-measured MSD and the simulations (Extended Data Fig. 6l, using as input parameter the average protrusiveness measured in late internalizing cells in MZ*lefty1/2* embryos - Extended Data Fig. 6j). This expanded window of competence for mesendoderm cell (un)jamming and autonomous internalization (Extended Data Fig. 6d-f') is consistent with the increased Nodal signalling activity, and mesendoderm cell protrusiveness, observed in MZ*lefty1/2* embryos as compared to stage-matched wt cells (Extended Data Fig. 6a, b, h-k). Finally, a prediction from our model is that while 50% epiboly stage donor cells are above the motility threshold for unjamming, they are expected to remain relatively close to the threshold. Thus, although nearly all cells internalize at the end of the simulation, they are predicted to do so with asynchronous dynamics (Extended Data Fig. 4g), a feature we also found in the experimental data (Extended Data Fig. 1c).

Numerical simulations of mesendoderm internalization with uniform cell-cell adhesion

As a minimal model for active adhesive tissues, we used a well-established framework of particle-based models[3–10], where each cell is represented by a point particle subject to interaction forces from its neighbours $F_{LJ}$, friction forces with its environment $F_f$ and exerting active migratory forces $F_a$. This type of model has been used in the past for understanding collective cell migration *in vitro*[3,5,9], as well as the theoretical signatures of active jamming in active soft matter[8,11]. It is well-suited for simulating zebrafish mesendoderm internalization movements, given that zebrafish mesendoderm is a tissue undergoing partial EMT with complex boundary conditions, which are difficult to implement in alternative models, such as Vertex or Voronoi models. As for the previous toy-model, which represented the dynamics of a cohesive donor cell cluster (modelled effectively as a single entity) by averaging the interaction with its surrounding $F_{LJ}$ in the form of a simple energy barrier, the system is heavily overdamped given the length and time scales considered, so that inertia is negligible and forces are balanced at all times for each particle.

The interaction force between two cells is described by the classical Lennard-Jones potential ("sticky spheres" with short-range repulsion and medium-range attraction), with an extended basin of attraction as in[10]. This allows cells to rearrange throughout migration while maintaining confluency. The potential from which the force derives ($F_{LJ} = dV_{LJ}/dr$) thus reads:

$$V_{LJ} = 4\epsilon(\frac{\sigma^{12}}{r^{12}} - \frac{\sigma^6}{r^6})$$

For $r \in \left[0, 2^{\frac{1}{6}}\sigma\right], V_{LJ} = -\epsilon$, for $\in \left[2^{\frac{1}{6}}\sigma, \bar{r}\right]$ *and* $V_{LJ} = 4\epsilon(\frac{\sigma^{12}}{(r-\bar{\sigma})^{12}} - \frac{\sigma^{6}}{(r-\bar{\sigma})^{6}})$ for $\in [\bar{r}, r_{cut}]$. Here, $\sigma$ is the characteristic cell size (which we use to set the length scale of the simulations, $\sigma = 1$), $\epsilon$ the scale of interaction forces, and $\bar{r} = 1.33, r_{cut} = 1.7$ to avoid cells from interacting with their next-nearest neighbours ($\bar{r} = 2^{1/6}\sigma + \bar{\sigma}$ from continuity). The frictional force of cells with the environment is classically modelled as a fluid friction $F_f = -\xi v$, where $v$ is the local cell velocity and $\xi$ a friction coefficient. We also added a small amount of Brownian noise to the simulations.

As we sought to model the internalization dynamics of the dorsal mesendoderm, we restricted ourselves to a two-dimensional (2D) description, which represents a sagittal cross-section of the dorsal marginal region of the zebrafish blastoderm. We thus simulated cells in a rectangular box of dimensions $L_x$ and $L_z$ ($x$ being the anterior-posterior axis and $z$ being the inside-out axis of the embryo). Based on our experimental observations of dorsal mesendoderm internalization at the blastoderm margin (Fig. 1b and Supplementary Video 1), we took $L_z = 7.5$ in our simulations and restricted ourselves to $L_x = 30$, as cells very far from the margin in the $x$ direction do not contribute significantly to the dynamics (Fig. 1a, b, Extended Data Fig. 1b and Supplementary Video 1). To initialize the simulations, we positioned 93 cells in a slab of three columns (from $z = 4$ to $z = 7$ and from $x = 0$ to $x_{margin} = 30$) together with small random noise in their initial positions. These cells are subject to three main forces: forces driving epiboly movements ($F^e$), active migration forces driving mesendoderm internalization ($F_a^i$) and anterior-directed active migration of internalized mesendoderm progenitors at the yolk syncytial layer (YSL) boundary ($F_a^a$; Fig. 1a', b and Supplementary Video 1)[12,13].

We then assigned each cell a specific Nodal signalling value, based on the nuclear accumulation of phosphorylated Smad2/3 complexes (pSmad2/3), a proxy for Nodal signalling activation[13–18], at the margin of 50% epiboly stage embryos (Fig. 2e and Extended Data Fig. 2h). The profile of nuclear accumulation of pSmad2/3 at the onset of gastrulation was well-fitted by a single-exponential decay of characteristic length scale $\lambda = 3.6 \pm 0.2$. Thus, we considered the Nodal signalling gradient as $N(x) = e^{(x-x_{margin})/\lambda}$. Note that we have normalized the maximal Nodal signalling value at the margin to 1, as this sets the scale for Nodal signalling in the simulations. Based on our experimental evidence (Fig. 2a-i, Extended Data Fig. 2a, b, g, 3c-i', k, l, 5a-f, g and 6a, b, h-k), the magnitude of active internalization forces $F_a$ (accessed by the rate and length of cell protrusions formed by mesendoderm cells) is proportional to Nodal signalling activity, whereas the directionality of these forces is not.

We assigned two components to active migration forces: internalization forces $F_a^i = -f^i N\, \vec{z}$ (directed towards the YSL and proportional to Nodal signalling) and anterior-directed migration forces $F_a^a = -f^a\, \vec{x}$ (constant for cells above a Nodal threshold $N_0 = 0.05$; Fig. 4a). Given that active anterior migration has been suggested to require contact with the YSL[19] and that mesendoderm cells closest to the YSL display faster anterior migration speeds (Extended Data Fig. 7c), we apply the forces $F_a^a$ to cells close to the YSL (defined as a range of $z \in [0, 1.5]$). We also made the simplifying assumption that all mesendoderm cells can migrate anteriorly ($f^a$ constant). However, relaxing this assumption, by assuming $f^a$ is proportional to Nodal signalling activity, did not yield significantly different internalization dynamics and still preserved positional information – although the overall anterior migration dynamics became expectedly slower (Extended Data Fig. 5n). *In vivo* active anterior migration forces are, at later stages of gastrulation, complemented by 3D convergence-extension movements[12], an effect we do not model here for the sake of simplicity.

Finally, we used reflecting boundary conditions so that cells cannot leave the simulation box and imposed a force on the most superficial cell layer to account for epiboly movements ($F^e = f^e\, \vec{x}$ if $z \in [L_z - 1, L_z]$). Importantly, incorporating this epiboly force, which can arise from the spreading of the overlaying enveloping layer (EVL) and/or directly from deep cell movements, was not strictly necessary and did not yield any significant changes to the results described here, as it simply translated cell movements by a constant speed. We nonetheless chose to incorporate them for a simpler visual comparison between simulations and data. Our experimental finding that cells had similar absolute speeds before (epiboly), during (internalization) and after (anterior migration) internalization (Extended Data Fig. 5h) led us to the simplifying choice of $f^e = f^a = f^i = 1$ (which we can set to 1 without loss of generality, as it sets the unit of forces in the simulations). Simulations were run for a total time $T_{tot} = 300$ (a "frame" of 150, for instance as shown in Fig. 4b represents a snapshot of a simulation at time 150) with background dissipation of $\xi = 500$, to match the experimentally-observed extent of mesendoderm internalization, whereby roughly 10-15 cell tiers undergo internalization (Extended Data Fig. 7c).

After parametrizing the simulations in this manner, we still had some free parameters, the most crucial being the interaction strength $\epsilon$. We thus performed simulations for a broad range of this parameter (see also subsection below) and found that it could be quite easily constrained, as too small values of $\epsilon$ (or high values of migration forces) resulted in the loss of tissue cohesion (Extended Data Fig. 5i, k), which we do not observe experimentally; whereas too large values of $\epsilon$ (or low values of migration forces) resulted in the tissue being jammed (Extended Data Fig. 5i, k). To quantitatively assess the sensitivity of the simulation predictions as a function of interaction strength $\epsilon$, we performed a parameter sweep and systematically calculated the fraction of simulations with loss of tissue cohesion and the $R^2$, as a metric of how well positional order is preserved (described in detail below). We found that tissue cohesion is only well-preserved for $\epsilon > 0.7$, while the $R^2$ started to decay below the values observed *in vivo* for $\epsilon > 1.0$. Thus, intermediate values of interaction strength in the range of $\epsilon = 0.7 - 1.0$ were able to reproduce well the experimental data (Extended Data Fig. 5i and Fig. 4b). In the simulations described in the main text (Fig. 4b, f, 5a and Supplementary Video 4), we used a value $\epsilon = 0.9$, corresponding to values of cell-cell adhesion on the same order as active migration forces.

<u>Simulation of mesendoderm internalization in wild type embryos</u>

One important feature of this numerical simulation is that it recapitulated the characteristic "turning" motion of mesendoderm cells at a single, well-defined location at the blastoderm margin (Fig. 4b and Supplementary Video 4), as seen *in vivo* (Fig. 1b and Supplementary Video 1). To quantitatively compare how positional order during mesendoderm internalization was preserved in the simulations and experimental data, we analysed the relationship between the relative positions of the first 7 cell tiers pre- and post-internalization, by recording their initial $x_i$ coordinate at the start of the simulation (N=20 simulations). Cell position was normalized between 0 and 1, so that $x_i = 0$ corresponds to the cell initially closest to the margin and $x_i = 1$ to the cell furthest away from the margin. We then recorded the relative positions $x_f$ post-internalization at the simulation time $t_0 = 150$, where the number of internalized cells was similar to the corresponding experimental time point (60 min, Fig. 1b-e and Extended Data Fig. 1b). If positional information in the simulations and experimental data were perfectly preserved, the first cells "in" should also be the first cells "out" of the marginal region and a perfect positive anti-correlation should be obtained ($y = 1 - x$, shown as a dashed line in Fig. 4c for example) between $x_i$ and $x_f$.

In simulations where the internalization force of mesendoderm cells scales with their Nodal signalling values (as described above), we found that, for intermediate values of interaction strength ($\epsilon = 0.9$), we could faithfully reproduce not only the experimentally observed correlation, but also the variance associated with it (Fig. 1f and 4c; see also statistical analysis section for the values of $R^2$ in each case). In contrast, performing simulations with constant internalization forces $F_a^i = -f^i \vec{z}$ for all mesendoderm cells ($N > N_0$, where $N_0 = 0.05$) revealed a markedly different scenario, where positional order was lost (Extended Data Fig. 5j). Importantly, performing simulations where all internalization forces were set to zero except for only a few leader cells ($f^i = 0$ for $N > 0.5$) still produced a highly ordered internalization process (Extended Data Fig. 5m), comparable to simulations where the internalization force scales with the Nodal signalling values and the wild type situation (Fig. 4b, c, Supplementary Video 1 and 4).

We also tested the impact of the magnitude of migration forces on the internalization process. To this end, we considered a situation where all cells experienced reduced migration forces compared to the previous simulations (both internalization and animal-pole directed $f^i$ and $f^a$). Reducing migration forces by a factor of 3 resulted not only in slower overall dynamics, but also impaired internalization movements, with simulations frequently showing tissues "stuck" in the initial stages of internalization (Extended Data Fig. 5k). This was due to all cells now behaving as "followers" with insufficient migration forces to drive the rearrangements necessary for their internalization. Taken together, these simulations are consistent with the sharp decay of autonomous internalization capacity observed in the transplantation assays (Fig. 1g-i and Supplementary Video 2) and further suggest that endowing a small fraction of leader cells with high internalization forces is necessary and sufficient to initiate ordered internalization movements.

Simulation of mesendoderm internalization in MZ*lefty1/2* embryos

To further challenge our model, we turned to MZ*lefty1/2* embryos, where the Nodal signalling gradient is expanded compared to wild type embryos[20] (Fig. 4d, e and Extended Data Fig. 6a, b). To this end, we first fitted the input Nodal gradient $N(x)$, based on the nuclear localization of pSmad2/3 complexes at the margin of 50% epiboly stage MZ*lefty1/2* embryos (Fig. 4d, e), and then tested whether this single parameter change could predict the experimentally observed mesendoderm internalization movements in these mutant embryos. In contrast to the wild type situation, the experimentally measured gradient of pSmad2/3 nuclear accumulation in MZ*lefty1/2* embryos could not be fitted by a single exponential; however a

Gaussian distribution $N(x) = N_0 e^{-\frac{(x-x_{margin})^2}{2\sigma_N^2}}$ provided a good fit, with $\sigma_N = 4.5$ and $N_0 = 1.15$ as best fit parameters.

Performing numerical simulations with this expanded Nodal signalling gradient (N=20 simulations) revealed strikingly different internalization dynamics compared to that of wild type embryos and simulations (Fig. 4f, b, 1b-f, Supplementary Video 4 and 1). Due to the expanded (Gaussian) shape of the gradient (Fig. 4d, e), many of the first cell tiers exhibited similarly high levels of Nodal signalling and thus internalization forces in the simulations. This resulted in the near-simultaneous internalization of several cell tiers, producing a "traffic jam" where many internalized mesendoderm progenitors try to initiate anterior migration (Fig. 4f and Supplementary Video 4). Furthermore, in these simulations, the cells initially closer to the margin were no longer positioned at the front of the internalizing tissue post-internalization, resulting in the loss of correlation between the positions of cells pre- and post-internalization (Fig. 4f, l and Supplementary Video 4).

Importantly, mesendoderm cell tracking in MZ*lefty1/2* embryos during tissue internalization revealed a close match to these simulations (Fig. 4g-l, Supplementary Video 4 and 5) and clear differences to the observations made in wild type simulations and embryos (Fig. 4b, c, Supplementary Video 4 and 1). In particular, the analysis of MZ*lefty1/2* embryos revealed that (i) the timing of mesendoderm cell internalization was neither any longer sequential, nor strictly correlated with initial cell position, as found in both wild type simulations and embryos (Fig. 4j, Extended Data Fig. 6c, Supplementary Video 5, Fig. 1e, Extended Data Fig. 1b, Supplementary Videos 1 and 4), leading to a weaker correlation as measured both by $R^2$ and slope (Fig. 4j and 1e), (ii) the distance travelled by mesendoderm cells in the $\vec{z}$ direction for the first hour of tissue internalization was nearly identical for the first 3-4 cells tiers (Extended Data Fig. 6c), instead of being largest for the first cell tier as found in wild type embryos (Extended Data Fig. 1b) and, (iii) comparison between the relative position of mesendoderm cells pre- and post-internalization showed no measurable correlation (Fig. 4k, l; while being tightly correlated in wild type simulations and embryos, Fig. 1f and Fig. 4c, see also statistical analysis section for the values of $R^2$ in each case). Collectively, this quantitative analysis of perturbation experiments supports the predictive power of our simulations and strengthens the conclusion that the shape of the Nodal signalling gradient regulates orderly initiation of tissue internalization, by controlling the fraction of leader and follower cells.

Influence of the range and dynamics of the Nodal signalling gradient on the internalization movements

To better understand the sensitivity of the model predictions in wild type and MZ*lefty1/2* embryos, we performed simulations by systematically increasing or decreasing the range of the Nodal signalling gradient, with the same exponential form as observed in wild type embryos $N(x) = e^{(x-x_{margin})/\lambda}$. We first ran simulations with larger length scales than observed in the experimental wild type data, taking $\lambda = 7$ cells (double the value fitted for wild type embryos). This resulted in highly disordered internalization movements (Extended Data Fig. 5o), similar to the simulations of MZ*lefty1/2* embryos with Gaussian gradients (see the previous section). In contrast to this, simulating a short-range Nodal signalling gradient ($\lambda = 1.75$ cells, half the value fitted for wild type embryos), did not show any strong difference to the wild type simulations, arguing that very few leader cells are sufficient to initiate tissue internalization, as long as follower mesendoderm cells still provide active anterior-directed migration forces (Extended Data Fig. 5p). This further supports the notion that a small fraction of leader cells, implemented by a fast-decaying Nodal signalling gradient, is highly efficient at initiating ordered internalization movements.

Finally, we also considered that the Nodal signalling gradient evolves in time. To this end, we updated the Nodal signalling value over time by a function $\Delta n(x)$ based on the local distance of a given cell to the margin, a well-known source of Nodal signals[15,21–24], with $\partial_t N(x) = \Delta n(x - x_{margin}(t)$. In these simulations, the margin was dynamically defined at each time point as the position of the most advanced cell $x_{max}$, and - as a simplifying assumption - we considered $\Delta n = \alpha e^{(x-x_{margin})/\lambda}$ (i.e. Nodal signalling gradient displaying the same decay distance as observed at 50% epiboly). This left a single free parameter $\alpha$ quantifying the rate of overall increase of Nodal signalling activity over time. For $\alpha = 0$, Nodal signalling activity in pre-internalizing progenitors decreases in time, as cells with higher Nodal signalling activity internalize. Conversely, for sufficiently large $\alpha$, Nodal signalling activity in pre-internalizing progenitors can increase in time. Since we found a progressive decrease in the nuclear accumulation of pSmad2/3 in pre-internalizing progenitors during gastrulation (Fig. 2e and Extended Data Fig. 2h), $\alpha$ is thus expected to be small. Exploring different physiological values for $\alpha$ revealed only small effects on the internalization process without affecting the overall tissue movements or the preservation of positional order (Extended Data Fig. 5q). In the initial simulations with the Nodal signalling gradient at 50% epiboly being set and constant, only cells with highest Nodal signalling values could internalize (Fig. 4b and Supplementary Video 4). In the simulations with an evolving Nodal signalling gradient, Nodal signalling in the follower cells increased once they reached positions close to the margin (i.e. when the first cells had already internalized), resulting in slightly faster overall internalization, but no qualitative change in order (Extended Data Fig. 5q).

Numerical simulations with non-uniform cell-cell adhesion

As discussed in the main text, our simulations suggested that gradients in Nodal signalling, and, consequently, cellular motility were sufficient to initiate ordered mesendoderm internalization, by specifying a small population of leader cells, which actively internalize and pull the remaining follower cells towards the inside of the blastoderm (Fig. 4b, c and Supplementary Video 4; also consistent with co-transplantation experiments of leader and follower cells, see Fig. 3d-f, Extended Data Fig. 4j, k and Supplementary Video 3). However, during later stages of these simulations, we observed a progressive loss of positional order (Fig. 5a, b and Supplementary Video 4), something we did not observe experimentally (Fig. 5b; see also statistical analysis section for the values of $R^2$ in each case).

In the simulations, this arose from our assumption of constant adhesion/interactions between all particles: the system is akin to a planar Couette flow rheometer, where opposing forces are exerted on the top versus bottom layers (epiboly forces towards the posterior/$\vec{x}$ axis and an active migration force towards the anterior/-$\vec{x}$ axis at the interface between internalized mesendoderm progenitors and the YSL). Thus, one expects constant gradients of velocity $v_x$ to be formed along the $\vec{z}$ axis, with highest velocities $v_x$ in internalized mesendoderm cells at the YSL boundary ($z \approx 0$), and zero velocities $v_x$ in internalized mesendoderm cells closer to the interface with non-internalized cells ($z \approx L_z/2$; Extended Data Fig. 7a for a schematic). Calculating the average velocity maps for the simulations considering uniform cell-cell adhesion confirmed this assumption, with mesendoderm velocities close to zero everywhere along the forming boundary between internalized and non-internalized cells and strong $v_x$ gradients along the $z$-axis irrespective of $x$ (N=20 simulations, Extended Data Fig. 7b). Such velocity gradient naturally resulted in a loss of positional information (Fig. 5a, b and Supplementary Video 4). In line with the better preservation of positional information observed *in vivo* (Fig. 5b, see also statistical analysis section for the values of $R^2$ in each case), we found that although such velocity gradient was also experimentally detectable near the margin[19], the velocity fields showed a clear non-zero velocity for mesendoderm cells further away from the margin (Extended Data Fig. 7c). This suggests that the adhesion/interaction between internalized and non-internalized cells is modulated in a position-dependent manner *in vivo*.

Based on this discrepancy between simulated and experimental data, we hypothesized that the Nodal signalling gradient could be used not only to set up the spatial pattern of cellular protrusive/motility forces, but also provide a graded code of relative adhesion/interaction strengths, which could, in turn, be used by leader cells to discriminate the correct followers to interact with and pull on. In particular, we theoretically explored the hypothesis that cells interact more strongly with cells at similar positions within the Nodal gradient and thus exhibit similar Nodal signalling activity (preferential/heterotypic adhesion, Fig. 5c and Extended Data Fig. 7d). Theoretically, such organization would be expected to be highly efficient in preserving positional information even at later stages of tissue internalization, since it allows cells to interact strongly with their immediate neighbours in the curvilinear axis of internalization and interact less with cells over the boundary between internalized and non-internalized cells (see Fig. 5c for a schematic). To implement this hypothesis in a minimal manner, we kept all parameters constant from the aforementioned wild type simulations, modifying only the Lennard-Jones potential with an interaction coefficient $\epsilon(N_i, N_j)$ between cells $i$ and $j$ that depends on their respective signalling values $N_i$ and $N_j$. The simplest theoretical assumption was to consider:

$$\epsilon(N_i, N_j) = \epsilon_0 e^{-\frac{(N_i - N_j)^2}{2\sigma_H^2}}$$

where the maximal adhesion/interaction strength $\epsilon_0$ (between cells with equal Nodal signalling values) is the same value as in previous simulations ($\epsilon_0 = 0.9$), and the interaction strength depends on the difference in Nodal signalling levels between contacting cells. The single parameter $\sigma_N$ quantifies how rapidly adhesion strength decays for a given Nodal signalling difference (large $\sigma_N$ means that differences in Nodal are inconsequential to cell-cell adhesion, while small $\sigma_N$ means that cell-cell adhesion is very sensitive to differences in Nodal signalling).

When performing a parameter-sweep for different values of $\sigma_H$, we found that $\sigma_H \gg 1$ expectedly gave rise to similar dynamics as observed in the previous simulations considering uniform adhesion (Fig. 4b, 5a, Extended Data Fig. 7b for $\sigma_H = 100$), whereas very small values of $\sigma_H < 0.3$ resulted in loss of tissue-level cohesion, since even small differences in Nodal signalling between cells along the anterior-posterior/x-axis are sufficient to locally lower cell-cell adhesion below the critical value required for cohesion. Intermediary levels of preferential adhesion ($\sigma_H = 0.6$), however, gave rise to internalization movements which closely resembled the experimental data (Fig. 5d and Supplementary Video 4), with good preservation of tissue-level cohesion and increasingly weak interactions at the boundary between internalized and non-internalized cells further away from the margin (Fig. 5d, Extended Data Fig. 7k and Supplementary Video 4). This resulted both in very similar average velocity maps for internalized mesendoderm cells and overall preservation of positional order in the simulated and experimental data (Extended Data Fig. 7c, k and Fig. 5e, see also statistical analysis section for the values of $R^2$ in each case; note that the time points in the simulations were chosen to match the average advance of the internalized tissue along the x-axis at each time point of the experimental data in Extended Data Fig. 7c). Indeed, when using $\sigma_H = 0.6$, strong gradients in velocity $v_x$ were only detectable close to the blastoderm margin at all time points, where Nodal signalling values between contacting cells remain rather similar. However, these gradients became less pronounced further away from the margin, as the difference in Nodal signalling values over the newly formed tissue boundary increases, thus reducing the interaction strength between internalized and non-internalized cells (see, for instance, cells at $x > 5$ in intermediate and late time points in Extended Data Fig. 7k).

This was also consistent with our co-transplantation experiments: whereas co-transplanting mesendoderm cells collected at the blastoderm margin from 50% epiboly and shield stage embryos typically resulted in cohesive clusters, co-transplantation of mesendoderm cells from 50% and 75% epiboly stage embryos did not (Fig. 3d-f, 5f-h, Extended Data Fig. 4j, k, Supplementary Videos 3 and 6). This behaviour is expected from the heterotypic/preferential adhesion model, given that the difference in Nodal signalling activity between mesendoderm cells from 50% and 75% epiboly stage embryos is bigger compared to the difference between cells from 50% and shield stage embryos (Fig. 2e and Extended Data Fig. 2h). Moreover, we also found that reducing the overall difference in Nodal signalling activity between leader and late follower cells, by co-transplanting wild type mesendoderm cells from 50% epiboly stage embryos and mesendoderm-induced cells from 75% epiboly stage embryos, partially rescued cohesion of the heterotypic clusters compared to co-transplantations of wild type mesendoderm cells collected from 50% and 75% epiboly stage embryos (Extended Data Fig. 7l-n, 3a-f, Fig. 5f-h and Supplementary Video 6).

This model with graded preferential adhesion bears resemblance to the hypothesis of heterotypic adhesion recently proposed to segregate ectoderm and mesoderm cells in *Xenopus* embryos[25]. In this case, ectoderm and mesoderm were considered as homogeneous populations, and heterotypic adhesion between these distinct germ layer progenitors was found to be disfavoured compared to homotypic interactions, namely ectoderm-ectoderm and mesoderm-mesoderm interactions. In our simulations, this would be equivalent to calculating the adhesion coefficient for cells with Nodal signalling levels set to either 0 (i.e. ectoderm) or 1 (i.e. mesoderm). However, running simulations under this assumption would result in the transition zone between mesoderm and ectoderm becoming highly fragile, since it corresponds to a very sharp boundary in Nodal signalling activity and would thus be characterized by weak heterotypic adhesion. Therefore, extending such binary heterotypic adhesion model to a preferential adhesion gradient, set by graded Nodal signalling, is not only more consistent with our experimental observations in co-transplantation experiments (Fig. 3d-f, 5f-h, Extended Data Fig. 4j, k, 7l-n, Supplementary Videos 3 and 6), but also constitutes a robust way of maintaining order while keeping tissue-level cohesion during complex 3D tissue remodelling.

Finally, we computationally explored alternative explanations for the observed experimental data by considering other functional forms of $\epsilon(N_i, N_j)$. Given that Nodal signalling has previously been found to increase cell-cell contact duration in prechordal plate cells[26], we explored the option that Nodal signalling-dependent differential adhesion might explain our experimental observations (Extended Data Fig. 7d, e). In a differential adhesion model[27–30], the interaction strength is defined as $\epsilon_i = \epsilon_0 + \alpha N_i$, and the classical assumption for mixtures of cells with different adhesion $\epsilon_i$ and $\epsilon_j$ is to consider the strength of adhesion between a given interacting pair equal to the geometric average of both interaction coefficients $\epsilon_{ij} = \sqrt{\epsilon_i \epsilon_j}$. In these simulations (considering the Nodal signalling gradient at the onset of gastrulation in wild type embryos), we found that internalized mesendoderm cells still showed strong interactions with non-internalized cells (Extended Data Fig. 7f, shown for $\epsilon_0 = 0.9, \alpha = 0.5$, all other parameters were kept as in Fig. 4b and 5a, d). This was clear both from average velocity maps (showing near-zero velocities of internalized mesendoderm cells near the interface to non-internalized cells even far away from the margin, see Extended Data Fig. 7g), as well as from the poor preservation of positional order (Extended Data Fig. 7h, i, see also statistical analysis section for the values of $R^2$ in each case). This is consistent with computational and experimental studies in *Xenopus*, suggesting that differential adhesion is not a robust mechanism to reduce cell mixing at tissue boundaries compared to heterotypic/preferential adhesion[25,28,29,31–33]. Collectively, these simulations argue against Nodal signalling-dependent differential adhesion being sufficient to explain the preservation of positional order during anterior migration of internalized mesendoderm progenitors.

This conclusion is also further supported by our triple co-transplantation experiments (Fig. 5f-h). Indeed, according with the differential adhesion hypothesis, cells collected at the blastoderm margin of 50% epiboly stage embryos would be expected to be the most adhesive cells, while cells collected from 75% epiboly stage embryos would be least adhesive cells ($\epsilon_{50\%} > \epsilon_{shield} > \epsilon_{75\%}$, Extended Data Fig. 7d). Consequently, contacts between cells collected from shield and 75% epiboly stage embryos should be weaker than those formed between cells collected from 50% and 75% epiboly stage embryos ($\epsilon_{50\%-75\%} > \epsilon_{shield-75\%}$, Extended Data Fig. 7d). This is, however, incompatible with our triple co-transplantation experiments, where we consistently observed $\epsilon_{50\%-75\%} < \epsilon_{shield-75\%}$ (Fig. 5f-h). The Nodal signalling-dependent preferential/heterotypic adhesion hypothesis instead predicts correctly this relationship ($\epsilon_{50\%-75\%} < \epsilon_{shield-75\%}$, Extended Data Fig. 7d), thereby allowing mesendoderm cells collected at the blastoderm margin of shield stage embryos to rescue the cohesion of these triple transplants, since it can establish relatively high adhesion with cells from both 50% and 75% epiboly stage embryos (Fig. 5f-h and see also Extended Data Fig. 7d for a schematic comparison of the predictions of the differential and heterotypic/preferential adhesion models).

Statistical analysis of the conservation of positional information during tissue internalization and average velocity maps

To analyse the conservation of positional information in the simulations during initial stages of tissue internalization (comparable to 0-60 min tracking in the experimental data of Fig. 1f and 4c, k, defined based on the number of internalized cells), we tracked the internalization movements of the first 7 tiers of cells (N=20 simulations; Fig. 4c, l and Extended Data Fig. 7h, j). We computed the relative position of cells along the X-axis (distance to the margin) before and after internalization as previously performed for the experimental data (see Methods for details). Similarly, to compute the tracking data for later stages of tissue internalization and anterior migration in the simulations (comparable to 60-90 min tracking data in the experimental data of Fig. 5b, e and Extended Data Fig. 7i, defined based on the number of internalized cells), we considered only internalized cells ($z < \frac{L_z}{2}$), which exhibit non-negligible Nodal signalling values ($N$>0.05). As we found experimentally that between 60 and 90 min, mesendoderm cells moved on average by 4 cell diameters, we defined, for each parameter set, the final time point of the simulations as the one for which cells have also moved by this amount (Fig. 5a, b, d, e and Extended Data Fig. 7f, i; see Methods for details).

To summarize the preservation of positional order between initial and final mesendoderm cell position, we calculated the coefficient of determination $R^2$ between initial and final position (when fitted by a function $y = 1 - x$ for 0-60 min and a function $y = x$ for internalized mesendoderm cells between 60-90 min). Thus, for the 0-60 min tracking, we calculated the coefficient of determination as $R^2 = 1 - S_{res}/S_{tot}$, where we defined the residuals $S_{res} = \sum_j \left(x_f^j - (1 - x_i^j)\right)^2$(where $x_f^j$ and $x_f^j$ are the initial and final normalized x-coordinates of cell *j*) and the total sum of square $S_{tot} = \sum_j \left(x_f^j - < x_f >\right)^2$. For the 60-90 min tracking, the definition was identical except $S_{res} = \sum_j \left(x_f^j - x_i^j\right)^2$ (function $y = x$ indicative of perfect correlation).

A perfect conservation of positional information would therefore translate to $R^2 = 1$, both for the experimental data and the simulations in all cases. Both dispersion around the ideal correlation slope ($y = 1 - x$ or $y = x$) and different slopes for the correlation between $x_f$ and $x_i$, would be expected to reduce the $R^2$ to values below 1. For the 0-60 min tracking, we found $R^2 = 0.63$ in wild type embryos (N=6 embryos) and $R^2 = 0.60$ in the corresponding simulations assuming uniform adhesion between all cells (Fig. 4c), showing a rather good quantitative agreement. Importantly, the simulations assuming preferential/heterotypic adhesion only marginally improved this initial correlation during the early stages of tissue internalization ($R^2 = 0.67$ in simulations with $\sigma_H = 0.6$, see Extended Data Fig. 7j), whereas the simulations assuming an adhesion gradient proportional to Nodal signalling (differential adhesion) worsened this correlation ($R^2 = 0.15$, see Extended Data Fig. 7h). Importantly, the correlation between initial and final position was fully lost in MZ*lefty1/2* embryos, where the experimental data showed a negative coefficient of determination $R^2 = -1.6$ (Fig. 4k, l). The simulations assuming an expanded Nodal signalling gradient closely reproduced this feature with a negative $R^2 = -1.1$ (Fig. 4l), reflecting the experimentally observed loss of positional order in MZ*lefty1/2* embryos.

For later stages of mesendoderm internalization and anterior migration (60-90 min tracking), we also calculated the $R^2$ for simulations assuming uniform cell-cell adhesion and found that it produced a poor correlation between initial and final mesendoderm cell position ($R^2 = 0.33$, see Fig. 5b), as did the simulations assuming Nodal signalling-dependent differential adhesion ($R^2 = 0.18$, see Extended Data Fig. 7i). On the other hand, simulations considering preferential adhesion ($\sigma_H = 0.6$) consistently predicted high correlations ($R^2 = 0.73$, Fig. 5e), which closely matched our experimental observations ($R^2 = 0.83$).

The analysis of the average velocity maps of internalized was performed similarly for the experimental data (Extended Data Fig. 7c) and the simulations with different model assumption (Extended Data Fig. 7b, g, k, N=20 simulations in each case; see Methods for details). As mentioned above, in the simulations, the analysis was restricted to internalized cells ($\boldsymbol{z} < \frac{\boldsymbol{L_z}}{\boldsymbol{2}}$) with non-negligible Nodal signalling levels ($\boldsymbol{N}$>0.05), in order to mirror the experimental tracking.
